# Caspase-1 self-terminates protease activity to enforce homeostasis and prevent inflammasome-driven diseases

**DOI:** 10.1101/2025.04.28.651119

**Authors:** Sabrina Sofia Burgener, Mark Thomas Milner, Shoumit Dey, Pooranee K. Morgan, Daniel G Blackmore, Emmanuelle Frampton, Gregory Miller, Monalisa Duarte de Oliveira, Kirsten Myra Kenney, Rinie Bajracharya, Ulrich Baumgartner, Thomas J.C Collins, Zherui Xiong, Quan Nguyen, Peter J. Meikle, Liviu-Gabriel Bodea, Andrew Clouston, Dave Boucher, Jürgen Götz, Andrew J. Murphy, Paul M Kaye, Kate Schroder

## Abstract

Signal shutdown mechanisms must exist to silence the potent inflammatory programs initiated by the caspase-1 (CASP1) protease, to allow inflammation to resolve and reinstate tissue homeostasis. It is unknown how CASP1 terminates its activity *in vivo*. Here, we use a knock-in mouse model in which the CASP1 CARD domain linker (CDL) is mutated to prevent self-cleavage (*Casp1.CDL* mice) to show that CASP1 CDL autoproteolysis terminates CASP1 activity *in vivo*. We examined these mice under homeostatic conditions and in response to major physiological challenges. In the brain, CASP1 CDL mutation caused anxiety-like behaviour under homeostatic conditions, and exacerbated hippocampal spatial learning deficits in the *APP23* genetic model of amyloid-induced neurodegeneration. In the bone marrow, CASP1 CDL mutation promoted steady-state granulopoiesis. In a model of diet-induced liver disease, CASP1 CDL mutation accelerated liver steatosis and promoted liver immune cell infiltration, inflammation and damage. In a liver healing model, CASP1 CDL mutation delayed disease resolution, indicating that CASP1 autocleavage is required to restore homeostasis after a major challenge to organ function. Our data reveal that CASP1 CDL self-cleavage terminates CASP1 inflammatory programs *in vivo* to maintain homeostasis in steady-state, restore homeostasis after a major challenge to organ function, and suppress inflammasome-driven diseases. These data identify CASP1 as a prime anti-inflammatory drug target, as CASP1 inhibitors may enforce homeostasis and prevent inflammasome-driven diseases.

## Introduction

Inflammasomes are multiprotein signalling hubs that assemble in response to disrupted homeostasis, including metabolic stress and the accumulation of protein aggregates, and provide an activation platform for the cysteine protease caspase-1 (CASP1). There are nine distinct human inflammasomes, several of which are linked to the pathogenesis of inflammatory and neurodegenerative diseases, and each of these inflammasome pathways converge to activate CASP1^1^. Thus, CASP1 is the central protease of inflammasomes, with active CASP1 cleaving pro-inflammatory cytokines IL-1β and IL-18 into their active, secreted forms to drive inflammation^2^. CASP1 also cleaves gasdermin D (GSDMD) to trigger a lytic form of cell death (pyroptosis) that causes tissue damage and further potentiates inflammation^1^.

The initiation and resolution of inflammatory pathways are tightly regulated. While mechanisms of inflammasome assembly and signalling are well studied, less is known of how this signalling complex is silenced. We identified a mechanism that controls CASP1 activation and deactivation within inflammasomes for cells cultured *in vitro,* in which CASP1 activity within inflammasomes is intrinsically self-limiting and controlled by a ‘proteolytic timer’ (**Suppl. Fig. 1A**). Here, CASP1 activation starts with the clustering of CASP1 monomers within the inflammasome complex, leading to proximity-induced dimerisation of the CASP1 protease domains, a necessary event for initiator caspases to gain proteolytic activity^2^. CASP1 p46 dimers then self-process at the interdomain linker (IDL) to unleash a fully active CASP1 p33/p10 species that cleaves substrates such as IL-1β. Terminating CASP1 activity requires a second self-cleavage event at the CARD-domain linker (CDL), which generates a p20/p10 species of dimeric CASP1; this species no longer contains the CARD domain that anchors CASP1 to the inflammasome to stabilise CASP1 dimers. Thus, p20/p10 dimers are ejected from the inflammasome and rapidly dissociate into inactive monomers, thereby terminating CASP1 protease activity^3^. These data indicate a model in which cellular CASP1 activation is coupled to its timely deactivation by CASP1 self-cleavage at the CARD domain linker (CDL)^3^. Such *in cellulo* investigations suggested that CASP1 CDL self-processing may silence CASP1 activity *in vivo*.

Mechanisms for inflammasome signal inhibition must exist to allow for tissue resolution following the inflammatory response to everyday homeostatic challenges, as well as major challenges that threaten organ function. Here, we generated a knock-in mouse model (*Casp1.CDL*) harbouring a compound point mutation at the major CASP1 CDL self-cleavage site (E102N/D103N), and examined several inflammasome-regulated tissue functions under homeostatic and challenge conditions. We show that CDL mutation within endogenous CASP1 dysregulates homeostasis in naïve mice, causing an anxiety-like phenotype and promoting steady-state bone marrow granulopoiesis. Further, suppression of CASP1 CDL cleavage accelerates disease progression in murine models of neurodegeneration and diet-induced liver disease. These data provide the first *in vivo* evidence that endogenous CASP1 self-regulates through a proteolytic timer mechanism to ensure timely CASP1 deactivation to enforce tissue homeostasis and suppress inflammasome-driven diseases.

## Results

### CASP1 self-cleavage at the CARD-domain linker (CDL) prevents hyperactive inflammasome signalling

CASP1 is the central protease in all human and mouse inflammasomes. CASP1 monomers cluster within the inflammasome to generate CASP1 p46 dimers that self-process at the IDL linker to generate fully active CASP1 p33/p10 species, while a second self-processing event at the CDL linker releases CASP1 p20/p10 species from the inflammasome to terminate CASP1 activity (**Suppl. Fig. 1A**)^3^. To understand how this mechanism silences the activity of endogenous CASP1 *in vivo*, we generated a knock-in mouse harbouring a compound point mutation in the major CDL self-cleavage site at position E102N/D103N (*Casp1.CDL*) (**Suppl. Fig. 1B-C**). To elucidate whether CASP1 CDL autocleavage at this site terminates endogenous CASP1 activity, we measured CASP1 cleavage of a fluorogenic synthetic substrate (Ac-WEHD-Afc) in WT and *Casp1.CDL* primary bone marrow macrophages (BMDMs). CASP1 was inactive in unstimulated and LPS-primed WT and *Casp1.CDL* BMDMs (**Fig. 1A**), indicating that CDL mutation did not confer CASP1 with constitutive activity in inflammasome-unstimulated cells. LPS-primed BMDMs were then stimulated with nigericin or ATP (to activate the NLRP3 inflammasome) or FlaTox (to activate the NLRC4 inflammasome), which triggered the accumulation of cleaved substrate over time in WT BMDMs. By contrast, CDL mutation to suppress CASP1 cleavage boosted CASP1 activity in both NLRP3- and NLRC4-activated *Casp1.CDL* BMDMs (**Fig 1B-D**), with peak CASP1 activity at 30 min (nigericin and ATP) and 60 min (FlaTox) post-stimulation. In keeping with CDL mutation conferring stimulus-induced CASP1 hyperactivity, *Casp1.CDL* BMDMs did not basally secrete IL-1β, and released significantly more IL-1β than WT BMDM when cells were exposed to nigericin, ATP or FlaTox (**Fig. 1E-G**). BMDMs produce TNF in a CASP1-independent manner, and accordingly, CDL mutation did not affect LPS-induced TNF release (**Fig. 1H-J**). CASP1 CDL mutation stabilised the active CASP1 p33/p10 species within BMDMs, leading to its enhanced capture by the biotin-VAD caspase activity probe (**Suppl. Fig 1D**). By suppressing p33/p10 autocleavage to p20/p10, CDL mutation blocked the generation of this inactive p20/p10 species that is not captured by the biotin-VAD caspase activity probe (**Suppl. Fig 1D**). Overall, these *in vitro* data confirmed that CASP1 CDL self-cleavage deactivates endogenous CASP1 to prevent stimulus-induced CASP1 hyperactivity, including upon the pro-IL-1β substrate.

**Figure 1.**
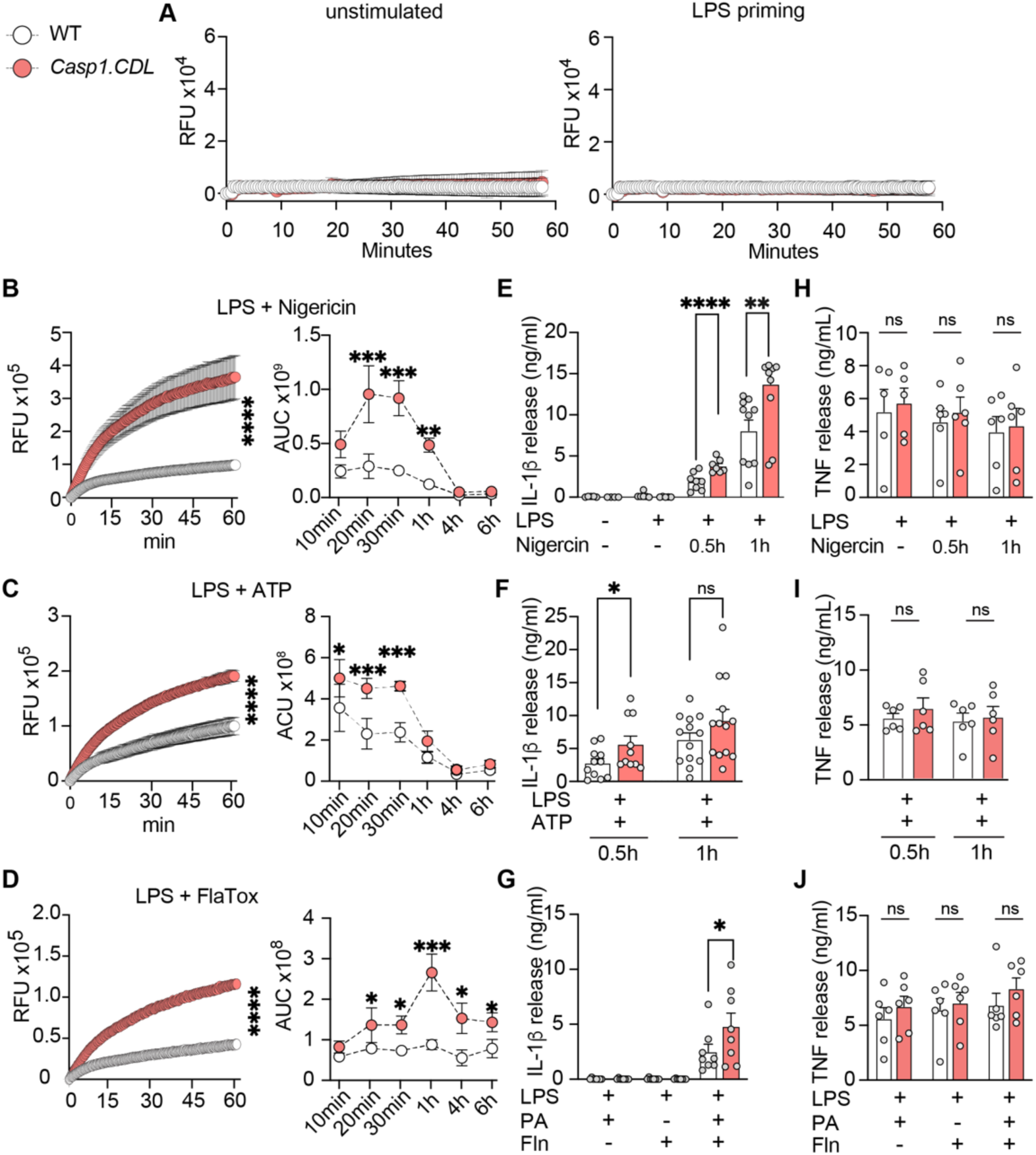
The CDL self-cleavage site of endogenous CASP1 suppresses CASP1 activities in inflammasome-activated macrophages. **a-i,** Bone marrow-derived macrophages (BMDMs) of WT or *Casp1.CDL* mice were either left untreated or primed for 4 h with 100 ng/ml ultrapure *E. coli* LPS (a) and stimulated with 5 µM nigericin or 2.5 mM ATP to induce NLRP3 signalling or **d+g+j**, primed for 4 h with 100 ng/ml ultrapure *E. coli* LPS and stimulated with 250 ng/ml PA and 125 ng/ml Fla1-LFn (FlaTox) to induce NLRC4 signalling. **a-d**, CASP1 activity was monitored in BMDMs that were untreated or LPS primed for 4 h (a), or LPS-primed BMDMs stimulated with nigericin (b), ATP (c) or FlaTox (d) for 10 min up to 6 h. At 10 min to 6h post-stimulation with inflammasome activators, the cell culture medium was replaced with caspase assay buffer containing the caspase-1 fluorogenic substrate, Ac-WEHD-Afc, and activity was measured for 60 min (see RFU kinetics; which relate to 30 min post-stimulation with nigericin or ATP, and 60 min post-stimulation with FlaTox). The area under the curve (AUC) was calculated for cells at each time point of inflammasome stimulation. Released mature IL-1β (**e-g**) or released TNF (**h-j**) was quantified by ELISA for LPS-primed macrophages stimulated for 30 and 60 min nigericin (e+h), 30 and 60 min ATP (f+i) and 60 min FlaTox (g+j). Dots represent BMDMs from individual mice (n=3-7 mice/genotype, assayed in 3-7 independent biological experiments), and data are shown as mean ± SEM. Data were verified for normality using a Shapiro-Wilk test, and analysed by two-way ANOVA using Šídák’s multiple testing correction. Statistical significance: * *p*≤0.05; ** *p*≤0.01, *** *p*≤0.001, **** *p*≤0.0001. See also Figure S1.

### *Casp1.CDL* mice exhibit anxiety-like behaviour at steady-state and spatial learning deficits in amyloid-induced disease

Having established that CASP1 CDL mutation causes signal-induced inflammasome hyperactivity *in vitro*, we then examined whether this occurs *in vivo* by examining steady-state brain functions. CASP1 deficiency in C57BL/6 mice decreases baseline and stress-induced anxiety-like behaviour during neurodevelopment^4,5^ and adulthood^6^. We thus investigated whether *Casp1.CDL* mutation affected mouse general behaviour, including ambulation, anxiety-like behaviour and learning. Independent of sex or genotype, mice exhibited decreased physical activity with age, reflected by reduced ambulatory distance and jump count and increased resting time with increased age (**Suppl. Fig 2**). *Casp1.CDL* female mice, but not male mice, showed a slightly increased body weight than WT mice at 9 and 12 months of age (**Suppl. Fig. 2F+L**). We next performed the elevated plus maze (EPM) behaviour test to examine anxiety-like behaviour and how this changed with aging. *Casp1.CDL* mice showed a marked increase in anxiety-like behaviour at 12 and 18 months of age, as compared to their WT counterparts – aged *Casp1.CDL* mice spent less time in the EPM open arms, more time in the closed arms, and less time moving around the maze as compared to WT (**Fig. 2A+B, Fig S3A-D**). To examine whether *Casp1.CDL* mutation affects cognitive function in mice, we next employed the Active Place Avoidance (APA) test of spatial learning and memory. *Casp1.CDL* mutation did not affect spatial learning in this paradigm at any age or in either sex (**Fig. 2C-D, Fig. S3E-H**). In all, these data indicate that while *Casp1.CDL* mutation does not affect the general physical activity levels of mice or their hippocampal spatial cognition in steady-state, it does increase their steady-state anxiety-like behaviour in both sexes.

**Figure 2.**
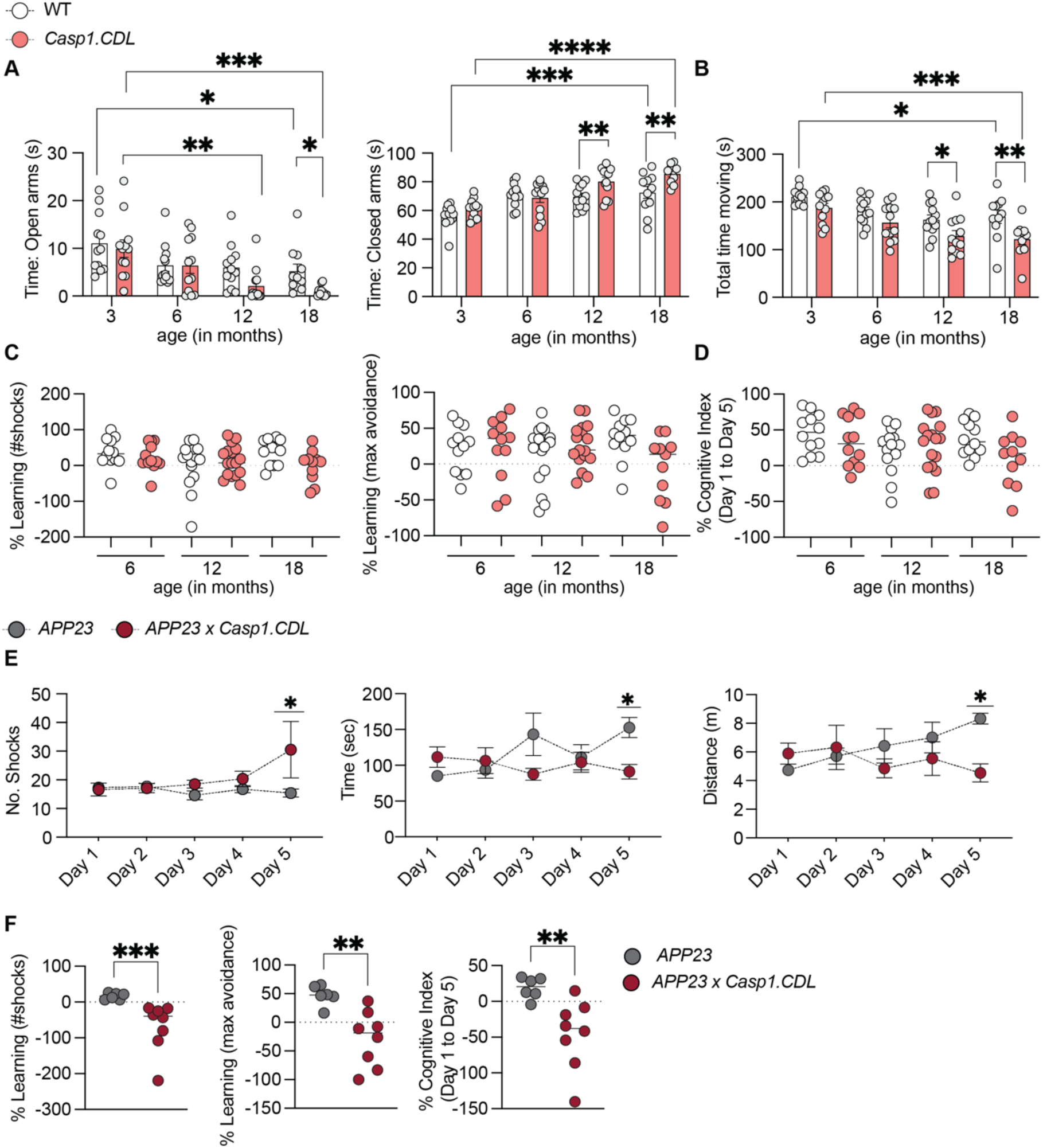
CASP1 CDL mutation drives anxiety-like behaviour during homeostasis and accelerates spatial learning deficits in amyloid-challenged mice. **a-d,** WT and *Casp1.CDL* male and female mice were aged and assessed for behaviour. **a-b**, Mice of 6-, 12- and 18 months were tested in the elevated plus maze (EPM), and assessed for time spent in open and closed arms (a) and movement over the duration of the test (b). **c-d**, Active place avoidance (APA) test was performed for 5 constitutive days with 6-, 12- and 18-month-old mice. % Learning between Day 1 and Day 5 was quantified, and shown as % learning based on shock number, % learning based on maximal time to avoid a shock (c) and a multi-parameter indicator of cognitive function (% cognitive index, d). **e-f**, 12-month-old *App23* and *App23.Casp1.CDL* male and female mice were tested using APA, as for panels C-D. **a-f**, Each dot represents individual mice (n=12-16 mice/group tested in 2-3 independent cohorts for a-d; n=6 *App23* and n=8 *App23.Casp1.CDL* mice, tested in 2 independent cohorts for e-f). All data are shown as mean ± SEM. Data for (a+b+e+f) were verified for normality using a Shapiro-Wilk test and analysed by two-way ANOVA with Šídák’s multiple testing correction. Non-parametric data were analysed using a Mann-Whitney U-test (f). Statistical significance: * *p*≤0.05; ** *p*≤0.01, *** *p*≤0.001, **** *p*≤0.0001. See also Figure S2-3.

Such findings from steady-state conditions prompted us to examine whether CASP1 CDL cleavage restrains disease parameters during a major challenge to brain function. We thus crossed the *Casp1.CDL* mice to the transgenic *APP23* mouse model that is well-characterized to cause deficits in spatial learning and memory. The *APP23* mice express the Swedish mutation (KM670/671NL) of the human APP gene (APP751 isoform) that causes familial Alzheimer’s Disease^7^, and are used to model neurodegenerative disease. Extracellular amyloid plaques are first detected at 6 months and increase with age, accumulating primarily in the cortex and hippocampus of *APP23* mice. APP23 mice exhibit deficits in APA-measured spatial learning by 3 months of age, and these deficits become increasingly severe with age^7,8^. Importantly for studies of CASP1, extracellular amyloid activates the NLRP3 inflammasome in murine microglia^9,10^, leading to CASP1-driven *in vivo* neuroinflammation and contributing to cognitive decline^11^. With the hypothesis that CASP1 CDL mutation leading to inflammasome hyperactivity would worsen amyloid-induced cognitive decline, we aged *APP23* and *APP23.Casp1.CDL* mice to 12 months of age and performed the APA paradigm. Indeed, *Casp1.CDL* mutation markedly exacerbated the spatial learning deficits of *APP23* mice, as indicated by *APP23.Casp1.CDL* mice experiencing a greater number of shocks over the 5- day learning period, due to their inability to learn to travel to actively avoid the shock zone (**Fig. 2E**). Multi-day analysis of shock number and maximum time to avoid the shock zone indicated that *APP23.Casp1.CDL* mice exhibited marked learning deficits as compared *APP23* mice. Multi-parameter data analyses provided an overall measure of cognition (cognitive index), which showed that *Casp1.CDL* mutation worsened the cognitive performance of *APP23* mice (**Fig. 2F**). Overall, these results demonstrate that CASP1 CDL mutation to suppress CASP1 inactivation magnified cognitive dysfunction and worsened spatial learning deficits in amyloid-induced disease. These findings position CASP1 as a central regulator of brain function in homeostasis and during a major challenge to tissue physiology.

### *Casp1.CDL* mice exhibit enhanced steady-state granulopoiesis

Our findings that CASP1 CDL mutation affected brain functions during homeostasis and amyloid-induced disease prompted us to examine the effects of this mutation in other organs. Inflammasome-dependent programs modulate immune cell birth, trafficking and lifespan. For example, IL-1β induces bone marrow granulopoiesis^12–15^, triggers neutrophil migration from bone marrow to tissues^12,16,17^ and delays neutrophil apoptosis^18–20^, while the pyroptotic signalling machinery can induce neutrophil death^21,22^. We thus investigated whether CASP1 hyperactivity caused by CDL mutation alters steady-state haematopoiesis by analysing the bone marrow and peripheral immune cell composition of *Casp1.CDL* mice. Relative to WT, *Casp1.CDL* mice exhibited a decrease in haematopoietic stem cells (HSC), particularly in short-term (precursor) multipotent HSC that sustain haematopoiesis, resulting in a reduction in granulocyte-monocyte progenitor cells (GMP) in the bone marrow (**Fig. 3A, Suppl. Fig. 4A-E**). Intriguingly, *Casp1.CDL* mutation was not associated with alterations in bone marrow monocyte abundance (**Suppl. Fig. 4B**) but did boost the frequency of circulating neutrophils in the blood and spleen (**Fig. 3B-C, Suppl. Fig. 4E**). Male *Casp1.CDL* mice showed a mild increase in white blood cell counts (WBC) in the blood (**Suppl. Fig. 4B**), while red blood cell (RBC) count, haematocrit and haemoglobin were unchanged. Reasoning that neutrophilia in *Casp1.CDL* mice could reflect a bias in colony formation and HSC differentiation towards neutrophils, we measured the capacity of HSCs to proliferate and differentiate *in vitro*. Both WT and *Casp1.CDL* mice showed a similar percentage of granulocyte-monocyte progenitor cells (GMP), while *Casp1.CDL* HSC exhibited lower potential to form myeloid-erythroid progenitors (MEP) in colony-forming unit assays (**Suppl Fig. 4F**). While this observation suggests a mild bias towards the myeloid lineage, it does not explain neutrophilia without concomitant monocyte elevation in *Casp1.CDL* mice. During infection or chronic situations of inflammation, IL-1β triggers G-CSF production to foster emergency granulopoiesis and ensure the rapid production of neutrophils. However, serum steady-state G-CSF levels were unchanged by CDL mutation (**Suppl Fig. 4G**). This indicated that CDL mutation did not induce granulopoiesis by promoting serum levels of G-CSF, the central regulator of neutrophil development and survival *in vivo*.

**Figure 3.**
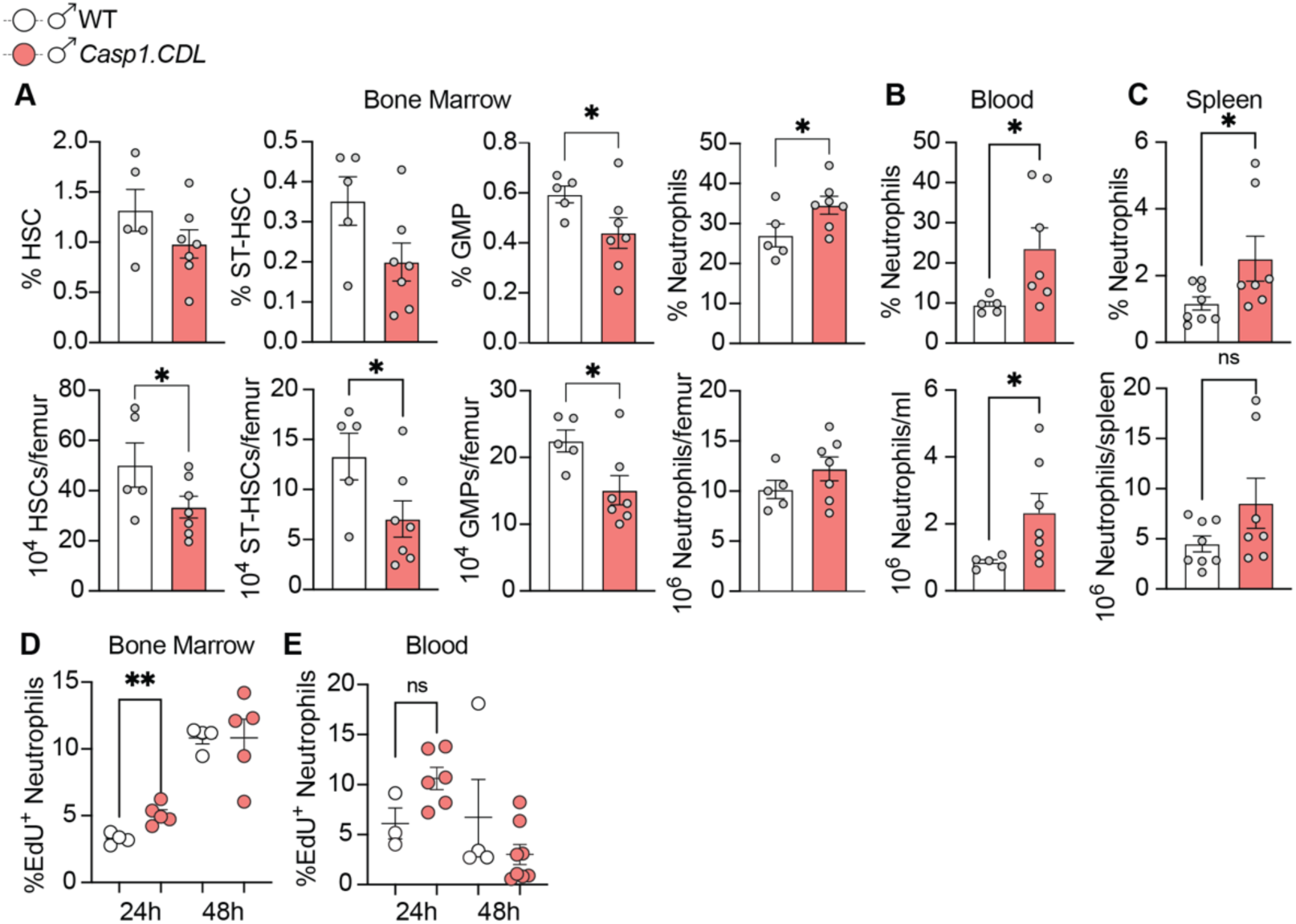
CASP1 CDL mutation promotes steady-state bone marrow granulopoiesis. 6-week-old male mice were analysed for percentage and abundance of: **a,** bone marrow hematopoietic stem cells (HSC), and mature neutrophils; **b,** circulating neutrophils; and **c,** splenic neutrophils. **d-e,** 6-week-old male mice were intraperitoneally injected with 2 mg/mL EdU and the percentage of EdU^+^ neutrophils in the bone marrow (d) and blood (e) was quantified 24 h and 48 h post-injection with EdU. **a-e,** Data are mean ± SEM, with dots representing individual mice (n=5-7 mice/genotype from 3-4 independent biological experiments in a-c; n=3-5 mice/genotype from 2 independent biological experiments in d-e). Data were assessed for normality using Shapiro-Wilk test. Parametric data were analysed by unpaired t-test (a-c) while non-parametric data were analysed by Mann-Whitney U-test (d-e). Statistical significance: * *p*≤0.05; ** *p*≤0.01. See also Figure S4.

Mature neutrophils are mobilised from the bone marrow into the blood upon cues such as injury or inflammation. To determine whether CASP1 CDL mutation affects neutrophil production and mobilisation, we pulsed mice with EdU to label newly-produced bone marrow neutrophils and track their egress from the bone marrow to the blood. 24 h after the EdU pulse, *Casp1.CDL* mice showed a higher percentage of EdU^+^ bone marrow neutrophils (**Fig. 3D)**, and a trend towards more EdU^+^ neutrophils in blood (**Fig. 3E**) compared to WT mice, without concomitant changes in EdU^+^ monocytes in either compartment (**Suppl. Fig. 4H**). These data indicate that *Casp1* CDL mutation promotes neutrophil production in the bone marrow, and perhaps their egress to the circulation, during steady-state hematopoiesis.

### Diet induces changes in hepatic lipid composition and MASLD

Our investigations into the steady-state behaviour (**Fig. 2**) and haematopoiesis (**Fig. 3**) of *Casp1.CDL* mice indicated that CASP1 CDL mutation perturbs these homeostatic features in the absence of a major organ challenge. Inflammasomes mediate sterile low-grade inflammation in conditions such as diet-induced liver disease, in which an unhealthy diet challenges homeostatic liver function. Here, a fat-rich diet triggers inflammasome-driven hepatic inflammation and resultant metabolic dysfunction-associated steatotic liver disease (MASLD) ^23^. MASLD encompasses a mild-to-severe disease spectrum, ranging from steatosis to inflammatory steatohepatitis (MASH) to liver fibrosis and eventually late-stage cirrhosis. MASLD progression to advanced fibrosis involves the swelling (ballooning) of lipid-loaded hepatocytes and their eventual death, which releases hepatocyte lipids into the liver microenvironment and activates hepatic innate immune cells such as liver-resident Kupffer cells, causing chronic liver inflammation and fibrosis^24,25^. To model MASLD progression *in vivo*, we employed a well-established model^23,26^ in which mice are fed a methionine-choline deficient (MCD) diet, relative to a control methionine-choline supplemented diet (MCS). We first assessed lipid accumulation in the liver after 2 weeks of MCD feeding, using lipidomics. Relative to their MCS-fed counterparts, MCD-fed WT mice showed elevations in the abundance of hepatic storage lipids such as Diacylglycerol (DG) and Triacylglycerol (TG) (**Fig. 4A, Suppl. Fig. 5A+B**); these are major constituents of lipid droplets, and their elevated abundance is an early sign of MASLD progression. MCD feeding also enriched liver Lyso-phosphatidylethanolamine (LPE), Cholesterol esters (CE) and Sphingolipids (Sph; specifically Sph such as Ceramides, Monohexosylceramide, Dihexosylceramides and Trihexosylcermide and Spingomyelin) (**Fig 4A, Suppl. Fig 5A**), lipids previously linked to hepatic inflammation, insulin resistance and oxidative stress^27,28^. We next examined key features of MASLD pathology. Relative to their MCS-fed counterparts, MCD-fed WT mice showed key histopathological features of MASLD, including liver steatosis (**Fig. 4B+C**), hepatocyte ballooning and inflammation (**Fig. 4C**) and an aggregate score of these features (non-alcoholic fatty liver disease (NAS) score, **Fig. 4C**). In all, these data indicate that MCD feeding alters the hepatic lipid profile to foster a lipid-laden hepatic environment that favours the initiation and progression of MASLD.

**Figure 4.**
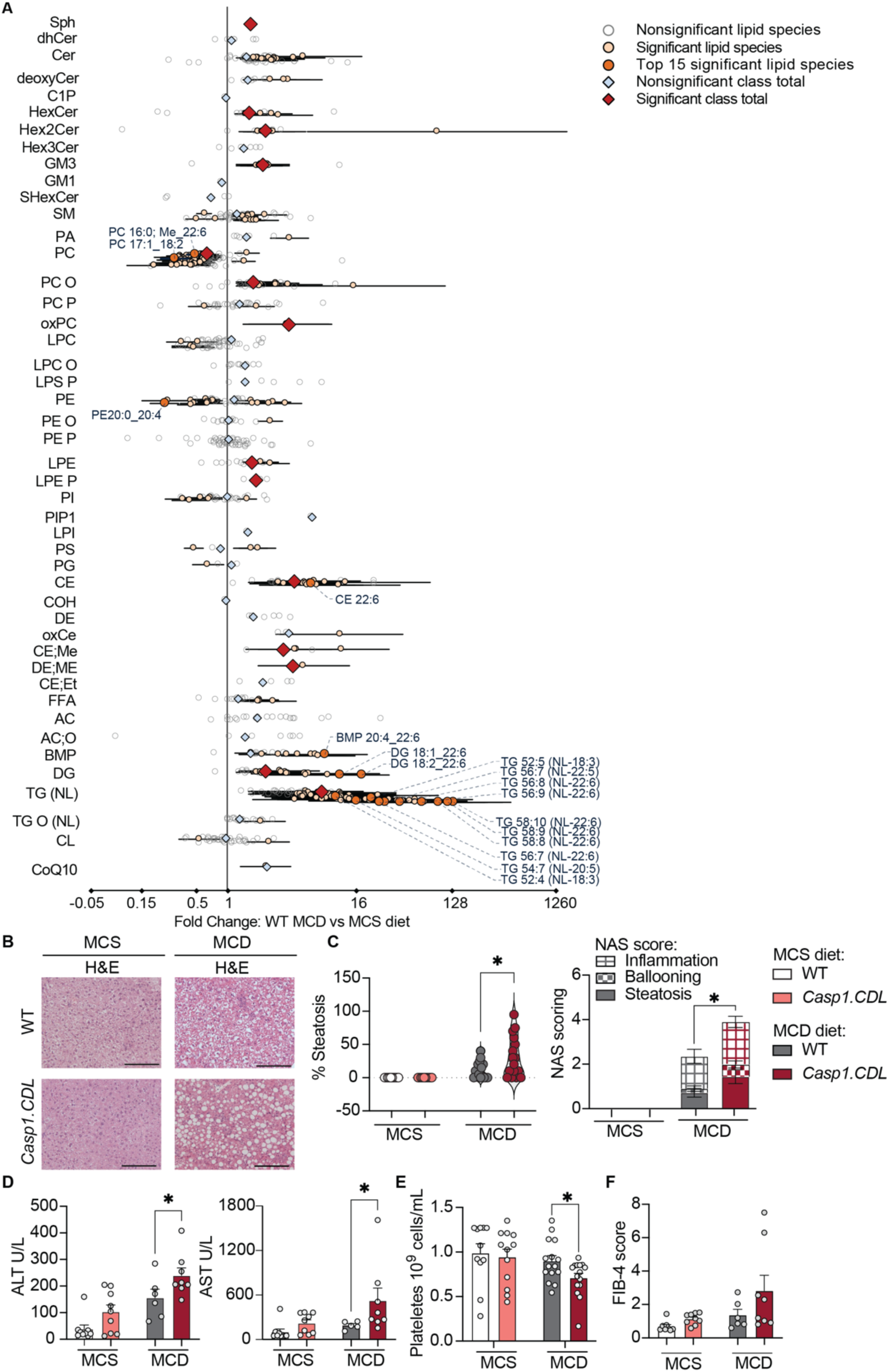
CASP1 CDL mutation exacerbates liver dysfunction during diet-induced liver challenge. 12-week-old male mice were fed a control methionine-choline supplemented diet (MCS), or a methionine-choline deficient diet (MCD) for 2 weeks to induce liver damage. **a**, Forest plot showing differences in the abundance of hepatic lipid species and classes in WT mice fed an MCD diet versus the control MCS diet. Data are shown as fold change, with upper and lower confidence intervals indicated. Statistically significant differences between MCD-fed and MCS-fed WT mice were determined using a paired t-test with FDR correction (n=8 for each diet). **b,** representative liver histology, shown by H&E staining (scale bar 100 μM). **c**, H&E-stained sections were blindly assessed for liver pathology according to percentage of steatosis, and scored using the NAS score (an additive score of steatosis, hepatocyte ballooning and inflammation). **d,** Serum levels of liver enzymes, alanine transaminase (ALT) and aspartate aminotransferase (AST), were measured as indicators of liver damage. **e,** The abundance of platelets in whole blood. **f,** FIB-4 score was calculated from ALT, AST and platelet counts. **c-f**, Dots in violin plots and scatterplots are data from individual mice (n=5-15 mice/group assessed in 4-5 independent experimental cohorts), except serum analyses, where two mice/group were pooled for analyte detection (d). Data are mean ± SEM, and were assessed for normality using Shapiro-Wilk test and analysed by a non-parametric Mann-Whitney U-test (c-f). Statistical significance: * *p*≤0.05; ** *p*≤0.01. See also Figure S5-8.

### *Casp1.CDL* mutation promotes early disease progression to inflammatory steatohepatitis

Specific lipid species activate inflammasome signalling *in vitro*^29–33^, and the MCD diet induces NLRP3 inflammasome signalling and resultant hepatic inflammation, leading to murine MASLD pathology *in vivo*^23^. We thus reasoned that while diet-induced changes in hepatic lipid profiles may be unaffected by inflammasome activity, hyperactive inflammasome signalling may exacerbate lipid-induced liver disease pathology in murine MASLD. To examine this hypothesis, we first explored lipid accumulation and liver pathology in mice fed the control MCS diet. *Casp1* CDL mutation did not affect the abundance of hepatic lipids in mice fed a control MCS diet (**Suppl. Fig. 6**), and neither WT nor *Casp1.CDL* mice fed the control MCS diet showed evidence of MASLD, including steatosis, hepatocyte ballooning or hepatic inflammation (**Fig. 4B+C**). We next investigated whether diet-induced changes in hepatic lipid profiles were affected by inflammasome activity. The MCD diet caused the accumulation of hepatic lipids in *Casp1.CDL* mice (**Suppl. Fig. 7+8**), similar to MCD feeding in WT mice (**Fig 4A**). In fact, while mouse diet induced a major impact on global lipid abundance, mouse genotype did not cause a generalised effect on lipid accumulation in mice fed either diet (**Suppl. Fig. 8A+B**), and this was also true of the proportion of individual lipid species (**Suppl. Fig. 8C-H**). Overall, these lipidomic analyses demonstrated that while the MCD diet triggered marked lipid accumulation in mice, hyperactive inflammasome signalling caused by *Casp1* CDL mutation did not alter hepatic lipids.

With the hypothesis that hyperactive inflammasome signalling will exacerbate lipid-induced liver disease pathology in murine MASLD, we examined whether CASP1 CDL mutation accelerates MASLD disease progression *in vivo.* Indeed, all MASLD-related pathological features including steatosis, hepatocyte ballooning, inflammation and NAS score, were elevated in *Casp1.CDL* mice relative to WT mice after two weeks of MCD diet feeding (**Fig. 4B-C**). Despite this short time frame, two weeks of the MCD diet in *Casp1.CDL* mice was sufficient to trigger substantial hepatocyte ballooning, a key histopathological indicator of MASH (**Fig. 4C**). Accordingly, MCD-fed *Casp1.CDL* mice showed elevated serum levels of liver enzymes such as alanine transaminase (ALT) and aspartate aminotransferase (AST), indicating more severe liver damage than the WT MCD-fed mice (**Fig. 4D**). The liver is an important source of the platelet growth factor thrombopoietin, and a drop in platelet count is an early diagnostic marker of liver damage that is commonly used to clinically assess MASLD. In keeping with CDL mutation worsening liver function, MCD-fed *Casp1.CDL* mice showed decreased platelets counts compared to MCD-fed WT mice (**Fig. 4E**). In the clinical diagnosis of MASLD, international guidelines recommend the Fibrosis-4 Index (FIB-4) as a first-line simple, non-invasive tool to estimate the risk of advanced liver fibrosis, with scores categorized into low, intermediate, or high-risk of fibrosis^34^. The MCD diet was associated with higher FIB-4 scores in both WT and *Casp1.CDL* mice compared to their MCS-fed counterparts (**Fig. 4F**). While differences between genotypes were not statistically significant, some *Casp1.CDL* mice had a score above 6 (**Fig. 4F**); this indicates an extremely high risk of fibrosis. In all, these data collectively suggest that CASP1 CDL mutation elevates several clinically-relevant parameters (e.g. steatosis, hepatocyte ballooning) and diagnostic markers (e.g. ALT, AST, platelet count) of diet-induced liver pathology. This in turn indicates that CASP1 CDL cleavage to terminate CASP1 activity is an essential step for restraining MASLD progression towards inflammatory steatohepatitis.

### CASP1 CDL mutation promotes liver inflammation to drive liver disease progression

We next sought to determine the mechanisms underpinning exacerbated disease progression in *Casp1.CDL* mice. To capture the heterogenous nature of disease pathology (**Fig. 4B+C**), we selected representative liver sections with mild (less than 20% steatosis), moderate (20-50% steatosis) and severe (above 50% steatosis) pathology from MCD-fed WT versus *Casp1.CDL* mice, alongside liver sections from MCS-fed controls, for analysis by spatial transcriptomics. Spatial gene expression analysis was performed for each 10x Genomics Visium spot (55µm resolution), and spots from 12 mice (4 groups; 3 mice per group) with a similar gene expression profile were integrated. Analyses identified six conserved clusters of spots that were present in all samples, regardless of the mouse diet or genotype (**Fig. 5A+B, Suppl. Fig. 9A-C**). The top 10 genes of each cluster were depicted for spots of the other five clusters. While cluster 3 spots showed a gene expression profile that overlapped with cluster 1, all other clusters showed distinct gene expression profiles (**Suppl. Fig. 9B**). Clusters 0 and 1 spots expressed genes that were consistent with parenchymal cells such as hepatocytes located within a hepatic region near the central (*Cyp2e1*) or portal vein (*Cyp2f*), respectively. Cluster 5 (*Hbb-bs*, *Hbb-a2*) was the least abundant spot cluster, and likely represents spots overlapping red blood cells. Visually, neither the diet nor the genotype affected the spatial distribution of the identified clusters, showing a heterogeneous distribution across the tissue sections (**Fig. 5B, Suppl. Fig. 9C**). Analysis of the proportional distribution of each spot cluster revealed a diet-dependent loss of cluster 0 spots in MCD-fed mice compared to MCS-fed mice, regardless of genotype (**Fig. 5C**). Compared to MCS-fed WT mice, MCD-fed WT mice showed decreased expression of genes involved in tryptophan and sulfur amino acid (SAA) metabolism (**Suppl. Fig. 9D**), which likely reflects the requirement for dietary methionine for the liver and other organs to produce SAA^35,36^.

**Figure 5.**
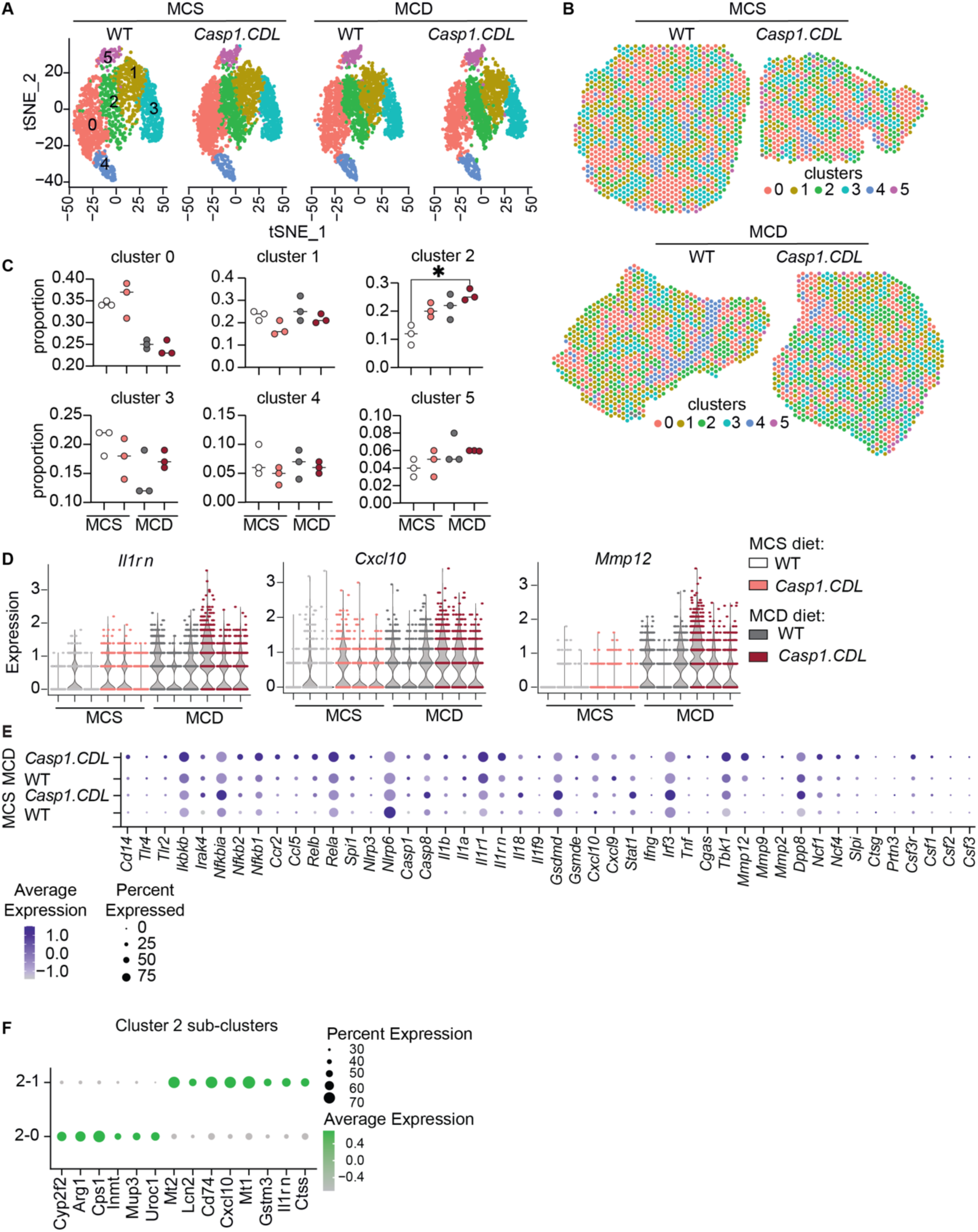
Visium spot clustering identified an immune-like signature increased by MCD feeding and CASP1 CDL mutation. Spatial mapping of transcriptomic clusters (visium spots) in mice fed a methionine-choline supplemented or deficient (MCS, MCD) diet for 2 weeks. 55μm diameter spots were clustered using principal components derived from their transcriptomic signatures (labelled as cluster 0-5), and visualised: **a,** in 2-dimensional t-SNE scatter plots; or **b,** according to their spatial x-y co-ordinates as on the underlying tissue (representative sections from MCS- and MCD-fed WT and *Casp1.CDL* mice). **c,** Each spot was assigned to a cluster, and the proportion of spots belonging to each cluster relative to all visium spots for all mice in an experimental group was calculated to give each cluster’s proportional distribution. **d,** Violin plot depicting the expression of *Il1rn, Cxcl10 and Mmp12* across all spots for each mouse. **e,** Bubble plot showing genes in cluster 2 in WT and *Casp1.CDL* MCS- or MCD-fed mice. The scale bar shows the average expression, and the size of the bubble represents the percentage of visium spots expressing the gene. **f,** Bubble plot showing sub-clustering of cluster 2 genes based on the top genes expressed in Cluster 2. **a-f,** Data are from n=3 mice per experimental group. **c**, Data are mean and individual mice (dots), and were analysed by non-parametric Kruskal-Wallis. Statistical significance: * *p*≤0.05. See also Figure S9.

Cluster 2 spots were increased in abundance by MCD diet and CASP1 CDL mutation (**Fig. 5C**). *Il1rn* was one of the most highly expressed genes of Cluster 2 spots (**Suppl. Fig. 9B**). In fact, across all mice and all spots, we observed a strong profile of immune/fibrotic-like transcripts, with *Il1rn*, *Cxcl10,* and *Mmp12* amongst the most highly expressed genes upregulated by MCD diet and CDL mutation (**Fig. 5D**). Given that the proportion of cluster 2 spots was enriched with MCD diet and CASP1 CDL mutation, we hypothesised that the immune-like signature typified by *Il1rn*, *Cxcl10,* and *Mmp12* profiles in global analyses may originate from cluster 2. Indeed, further analyses of the genes expressed within cluster 2 spots showed a strong correlation with innate immune cell activation pathways linked to NF-*κ*B, Toll-like receptor and IL-1R signalling, with elevated expression of *Ikbkb*, *Rela*, *Il1r1*, *Il1rn*, *Tbk1* and *Mmp12* in MCD-fed *Casp1.CDL* mice compared to MCD-fed WT mice, and with low or no expression in livers of mice fed the MCS diet (**Fig. 5E**). Gene enrichment pathway analysis to re-cluster the genes expressed within cluster 2 spots identified a distinct gene expression profile of two sub-clusters (2-0 and 2-1) therein (**Fig. 5F**), indicative of distinct sub-niches within visium spots. Sub-cluster 2-1 expressed elevated levels of *Il1rn* and *Cxcl10,* while sub-cluster 2-0 expressed genes related to parenchyma (**Fig. 5F**). Thus, cluster 2 visium spots are likely to represent immune cells in close communication with liver parenchymal cells. Collectively, these analyses indicate that cells captured by cluster 2 spots exhibit an immune-like signature that was most abundant with the MCD diet and increased further in *Casp1.CDL* mice, and may be responsible for driving exacerbated MCD-induced liver disease in *Casp1.CDL* mice.

Upon IL-1β binding to IL-1R, a cascade of downstream signalling (e.g. activation of NF-*κ*Β), gene regulation (e.g. upregulation of *Il1rn)* and cell activation occurs, ultimately resulting in collagen deposits, fibroblast activation, and immune cell activation within the liver^37–39^. Upregulation of the NF-*κ*Β target gene, *Il1rn*, by MCD diet and CASP1 CDL mutation as indicated by global and cluster 2 analyses (**Fig. 5D+E**), alongside the upregulated expression of the IL-1 signalling machinery (e.g. *Il1r1*, *Ikbkb*, *Rela*, *Il1r1*, *Tbk1;* **Fig 5E**), suggested that MCD diet and CASP1 CDL mutation may sensitise cells within cluster 2 spots to cell signalling by IL-1β:IL-1R. We thus sought to determine which spots express IL-1R, and which spots upregulate IL-1R expression with MCD diet and CASP1 CDL mutation. Spatial gene expression analysis for *Il1r1* for all visium spots (regardless of cluster) showed that *Il1r1* expression was strikingly enhanced by CASP1 CDL mutation, and was further increased when mice were fed the MCD diet (**Fig. 6A**, **Fig. 6B** all spots panel). The global trend for *Il1r1* upregulation by MCD diet was also evident when analyses were confined to cluster 0 and 2 spots (**Fig. 6B**). However, CASP1 CDL mutation upregulated *Il1r1* in cluster 2 but not cluster 0 spots in MCD-fed mice (**Fig. 6B**). Collectively, these analyses suggest that cluster 2 cells are poised to sensitively respond to IL-1β, and indeed may be generating IL-1-driven signalling outputs (e.g. *Il1rn*) in response to MCD diet and CASP1 CDL mutation. The striking loss of cluster 0 spots (hepatocyte signature with *Il1r1* upregulated by MCD diet; **Fig 5C, 6B**) in mice fed an MCD diet is consistent with lipid-induced hepatocyte death. Given that IL-1 signalling exacerbates lipid-induced hepatocyte death ^37^ and triggers myeloid cell recruitment ^40–42^, we next sought to determine whether hyperactive inflammasome signalling affects the cell composition or cell spatial organisation in the liver.

**Figure 6.**
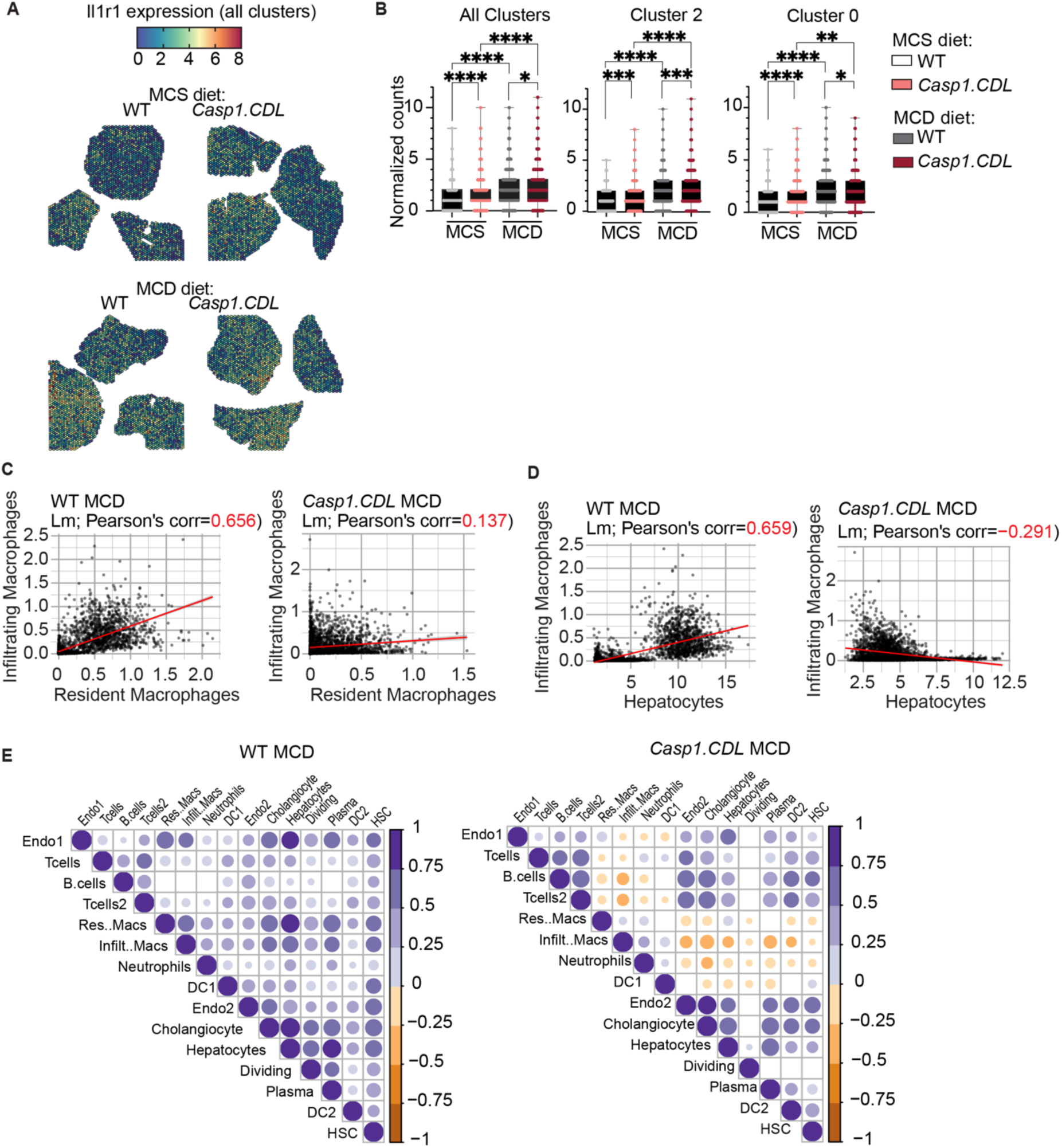
CASP1 CDL mutation uncouples the spatial co-location of infiltrating macrophages, resident macrophages and hepatocytes. **a,** Spatial gene expression of *Il1r1* abundance across all spots for livers of each mouse. **b,** *Il1r1* abundance in all spots, as well as spots belonging to cluster 2 or cluster 0. **c-d,** Cell2location predicted linear relationships of cellular abundance to approximate the relationships (Pearson’s product-moment correlation) of resident versus infiltrating macrophages (c), or hepatocytes versus infiltrating macrophages (d), as re-analysed from GSE129516 in visium spots. **e,** Pairwise Pearson’s correlations were represented as a correlation plot between predicted linear cell type relationships, to infer spatial co-location in the MCD-fed mouse groups. **b,** Data are represented as scatter plots for n=3 mice/group from 3 independent experiments, and were analysed using Kruskal-Wallis non-parametric mean rank between groups. Statistical significance: * *p*≤0.05; ** *p*≤0.01, *** *p*≤0.001, **** *p*≤0.0001. See also Figure S10.

### CASP1 CDL mutation uncouples the spatial co-location of infiltrating macrophages, resident macrophages and hepatocytes

We next investigated how liver cellularity may be altered by MCD diet and CASP1 CDL mutation to favour an immune-like gene signature, by further characterising the cellular composition of cluster 2 spots. We performed cellular deconvolution of all visium spots using a published scRNAseq dataset that profiled diet-induced murine liver disease^43^. Our earlier analyses of strongly expressed transcripts suggested that cluster 2 spots likely captured immune cells and hepatocytes (**Fig 5F**). Deconvolution analyses confirmed that cluster 2 spots captured a mixture of cell types including hepatocytes, hepatic stellate cells and several immune cells (**Suppl. Fig. 10A+B**).

Given that CDL mutation boosts CASP1-driven inflammatory programs (**Fig. 5E**), we hypothesised that this mutation will accelerate the local inflammatory response triggered by hepatic lipid accumulation and hepatocyte ballooning (**Fig. 4B-C**). Lipid-induced hepatocyte death releases danger-associated molecular patterns (e.g. ATP) and hepatocyte lipids that are likely to activate inflammasome signalling in liver-resident macrophages, fostering an IL-1 signature that causes immune cell infiltration into the liver, and boosts hepatocyte death in a feed-forward loop. By assessing cellularity, we determined that cluster 2 spots capture resident macrophages (Kupffer cells), infiltrating macrophages and neutrophils (**Suppl. Fig. 10B+C**). We next examined the spatial distribution of cluster 2 visium spots, and observed that resident macrophages (Kupffer cells) and infiltrating macrophages were distinctly spatially arranged in MCD-fed WT versus *Casp1.CDL* mice (**Suppl. Fig. 10C**). To quantify the spatial colocation of individual cell types, we modelled the proximity of one cell type to another by correlating the abundance of each cell type to another using Pearson’s product-moment correlation (**Fig. 6C-E**). Resident macrophages were strongly correlated with infiltrating macrophages in MCD-fed WT mice (r=0.66), suggesting these cell types are co-located (**Fig. 6C+E**). Intriguingly, MCD-fed *Casp1.CDL* mice showed a trend towards more infiltrating macrophages and a loss of resident macrophages, and the location of these two cell types was less strongly correlated (r=0.137) than in MCD-fed WT mice (**Fig. 6C**). Histological analyses using CD68 as a marker of resident and infiltrating macrophages confirmed that MCD feeding elevated the bulk abundance of macrophages in the liver (**Suppl. Fig. 10D-F**). Infiltrating macrophages have distinct morphology (round and compact; **Suppl. Fig. 10D – orange inserts**) compared to resident macrophages (elongated; **Suppl. Fig. 10D – purple inserts**). Infiltrating macrophages were noticeably more abundant in *Casp1.CDL* versus WT MCD-fed mice (**Suppl. Fig. 10D-F**). The MCD diet also induced the infiltration of neutrophils (Ly6G+) into the liver (**Suppl. Fig. 10H-K**). Macrophage and neutrophil infiltration were evident after 2 weeks of the MCD diet (**Suppl. Fig. 10D+E, H+I**), and also during advanced MASLD (4 weeks of MCD feeding, **Suppl. Fig. 10F+G, J+K**). We next examined the proximity of infiltrating macrophages to hepatocytes using spatial transcriptomic analyses, and observed that infiltrating macrophages were positively correlated with hepatocytes in MCD-fed WT mice (r=0.659) but negatively correlated in MCD-fed *Casp1.CDL* mice (r=-0.291) (**Fig. 6D**). To gain a broader view, all cell types captured by cluster 2 spots were analysed for co-location with one another within a spot. Amongst all these cell-cell relationships in MCD-fed mice, the CASP1 CDL mutation-driven spatial uncoupling of myeloid cell types with one another and with hepatocytes was most striking (**Fig. 6E**). These data suggest that infiltrating macrophages are proximal to resident macrophages and hepatocytes in WT MCD-fed mice, and inflammasome hyperactivity uncouples these spatial arrangements and the cell-cell communications they enable in *Casp1.CDL* mice.

### The CASP1 CDL is required for liver healing and restoring homeostasis

Our previous investigations into major physiological challenges to brain (**Fig. 2**) and liver (**Fig. 4-6**) function demonstrated that CASP1 CDL mutation worsened disease outcomes. It remained unclear whether CASP1 CDL cleavage is required to reinstate homeostasis after the removal of a major organ challenge. To address this, we employed a liver healing model in which we fed mice the MCD diet for 4 weeks to induce inflammation, liver damage and advanced MASLD (**Suppl. Fig. 11**) and then switched their diet to the healthy control MCS diet to monitor liver recovery (MCD-MCS). The liver is a remarkable organ, as it can rapidly repair itself by creating new tissue to replace damaged cells. This unusual feature of liver biology makes it an ideal organ to study the molecular and cellular mechanisms that restore homeostasis and organ function after a major challenge. The regenerative capacity of the liver is also important clinically; in the absence of disease-modifying therapies, clinical strategies for treating diet-induced liver have traditionally relied upon recommending lifestyle interventions (e.g. switching to a healthy diet) to induce liver healing and restore liver function.

Our previous studies showed that *Casp1.CDL* mice exhibited more rapid pathology than WT mice after 2 weeks of MCD feeding (**Fig. 4**). Over a further two weeks of MCD feeding, the pathology of the *Casp1.CDL* mice plateaued while WT continued to climb, with the outcome that after 4 weeks MCD feeding to model advanced MASLD, disease pathology was similar between these genotypes (**Suppl. Fig. 11**). After 4 weeks MCD feeding, we then switched mice to the control (MCS) diet and assessed liver healing (MCD-MCS). WT mice demonstrated a remarkable and almost complete disease recovery (**Fig. 7A-F**), with markers of steatosis, body weight and liver damage returning to baseline within a week of being on a healthy diet (MCD-MCS versus MCS, **Fig. 7A-F**). Given that MCD feeding changes the hepatic lipid content (**Fig. 4A**), we next assessed whether MCS feeding for a week to allow liver healing restored normal hepatic lipid abundance after MCD-induced injury. While some hepatic lipids remained perturbed by prior MCD feeding (**Supp Fig. 12A-B**), these perturbations were much less striking than MCD-alone feeding (**Fig. 4A**), indicating that WT mice fed MCD-MCS mice undergo recovery following MCD-induced MASLD.

**Figure 7.**
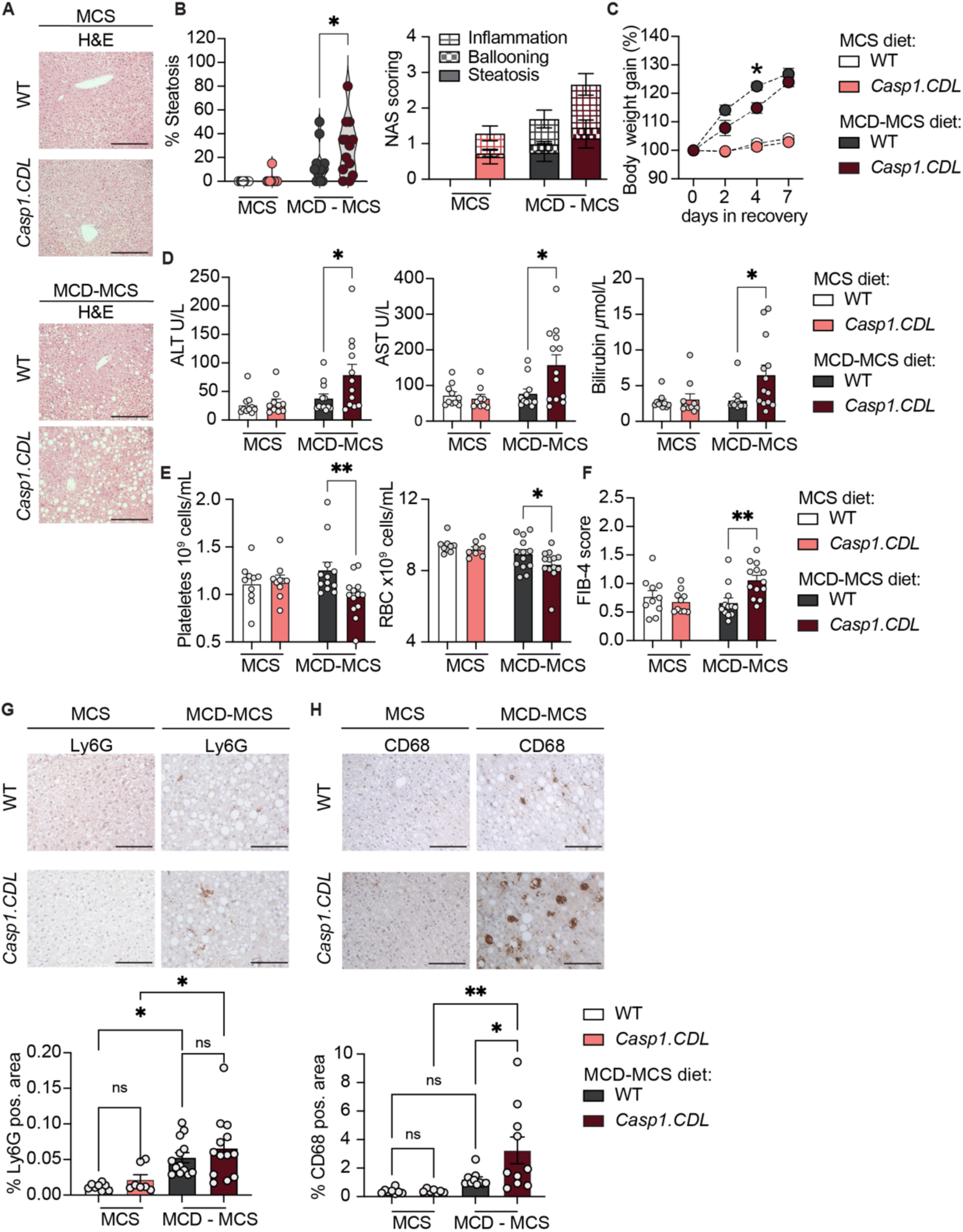
The CASP1 CDL promotes inflammatory resolution during liver healing. 12-week-old male mice were MCD-fed for 4 weeks to induce liver damage, and then switched to the control MCS diet for 7 days to allow liver healing (MCD-MCS), and compared to mice fed an MCS diet for 5 weeks. **a,** representative H&E staining of liver slices (scale bar 100 μM). **b**, H&E-stained sections were blindly assessed for liver pathology according to the percentage of steatosis, and scored using the NAS score (an additive score of steatosis, hepatocyte ballooning and inflammation). **c,** body weight gain following the switch in diet. **d,** Serum levels of liver enzymes, alanine transaminase (ALT), aspartate aminotransferase (AST) and bilirubin were measured as indicators of liver damage. **e,** The abundance of platelets and red blood cells (RBC) in whole blood. **f,** FIB-4 score was calculated from ALT, AST and platelet counts. **g**, Livers were stained using DAB immunohistochemistry for neutrophils (anti-Ly6G) and macrophages (Kupffer cells + infiltrating macrophage; anti-CD68). CD68 and Ly6G positive area was quantified using Fiji (ImageJ) analysis of the average signal of three liver pieces per mouse. Scale bar 100 µm. **b-g**, Data are mean ± SEM and violin plots and scatterplots show data of individual mice n=7-15 mice/group from 3-4 independent experiments, except serum analyses, where two mice/group were pooled for analyte detection (d). Data were assessed for normality using the Shapiro-Wilk test. Parametric data was analysed by two-way ANOVA (b) or one-way ANOVA (c+g) and non-parametric data was analysed by Mann-Whitney test (d, e, f). Statistical significance: * *p*≤0.05; ** *p*≤0.01, *** *p*≤0.01. See also Figure S11-12.

CASP1 CDL mutation prevented MASLD recovery. MCD-MCS-fed *Casp1.CDL* mice exhibited more severe liver steatosis (**Fig. 7A+B**), slower weight gain (**Fig. 7C**), ongoing liver damage indicated by higher serum levels of ALT, AST and bilirubin (**Fig. 7D**), and decreased platelet and red blood cell counts (**Fig. 7E**) compared to MCD-MCS-fed WT mice. Their elevated FIB-4 score (**Fig. 7F**) indicated that *Casp1.CDL* mice remained at a higher risk of fibrosis than WT mice. We next examined whether CASP1 CDL mutation suppressed the normalisation of hepatic lipids when mice were switched to the MCS diet. In keeping with lipids being upstream of inflammasome signalling, CASP1 CDL mutation did not affect lipid abundance in MCD-MCS-fed mice (**Suppl. Fig 13**). Both MCD-MCS-fed WT and *Casp1.CDL* mice showed a partial restoration of their overall lipid profile towards the baseline control MCS-fed mice (**Suppl. Fig. 14**) with the proportions of individual lipid species being generally comparable in MCD-MCS-fed WT mice compared to MCD-MCS-fed *Casp1.CDL* mice. Thus, while mouse diet dramatically altered the global lipid profile, mouse genotype did not affect overall hepatic lipid abundance, and induced only minor variations in the proportion of some individual lipids.

Another key feature associated with clinical MASLD mirrored in MCD-induced pathology is myeloid cell influx into the liver. MCD feeding for 2 or 4 weeks induced macrophage and neutrophil infiltration into the livers of both WT and *Casp1.CDL* mice, with bulk myeloid abundance largely unchanged by mouse genotype (**Supp Fig. 10D-K**). Here, myeloid cell infiltration is indicative of the inflammatory liver environment fostered by hepatic lipid accumulation during MCD feeding. To determine whether inflammasome hyperactivity in *Casp1.CDL* mice delays the resolution of inflammation in the liver, we assessed myeloid cell abundance in the livers of MCD-MCS-fed *Casp1.CDL* mice versus WT mice. MCD-MCS-fed mice exhibited greater neutrophil abundance compared to MCS-fed littermates, however, this was unaffected by CASP1 CDL mutation at this time point (**Fig 7G**). Most CD68-positive macrophages in MCD-MCS-fed WT mice were elongated, similar to the resident macrophages of the MCS-fed mice (**Fig. 7H**). By contrast, the round morphology of CD68-positive macrophages in MCD-MCS-fed *Casp1.CDL* mice indicated the sustained presence of liver-infiltrating macrophages (**Fig. 7H**). CASP1 CDL mutation also boosted the abundance of bulk macrophages in MCD-MCS mice, confirming that CASP1 CDL mutation sustained the presence of infiltrating macrophages (**Fig. 7H**). These data collectively indicate that inflammasome hyperactivity, caused by suppressed CASP1 CDL cleavage, promotes inflammation and blocks liver healing after a major, diet-induced challenge to liver homeostasis.

## Discussion

A central tenet of immunology is that potential threats to organismal homeostasis trigger inflammatory programs to eliminate the threat, after which inflammation is silenced to restore homeostasis and normal tissue functions. We understand how inflammatory programs are silenced *in vivo* for many innate immune pathways (e.g. toll-like receptors, cytokine receptors, and common signalling modules such as those activating NF-*κ*B or p38 MAPK). Here, switching off inflammatory programs generally involves transcriptional programs of signal feedback inhibition. For example, IL-1 signalling via the IL-1R triggers a signalling cascade that activates the NF-*κ*B transcription factor and its ensuing inflammatory programs, while NF-*κ*B also induces the expression of the NF-*κ*B inhibitor, I-*κ*B, and the IL1-R antagonist (encoded by the *Il1rn* gene) to then feedback to silence inflammatory signalling by NF-*κ*B and the IL-1R. Inflammasomes trigger potent inflammatory responses by regulating proteolysis, rather than transcription. Unlike transcriptional changes, which are inherently reversible, protein cleavage is not reversible. It has remained a major challenge to identify fundamental mechanisms that silence proteolytic signalling cascades and their resultant biological programs, such as inflammasome-driven inflammation.

Inflammasome-driven tissue responses to infection and injury are transient. In settings in which inflammasome-driven programs eliminate infection, inflammasome signalling must then be switched off to prevent chronic inflammation and disease. Most inflammasomes activate the protease, CASP1, which elicits inflammation by cleaving pro-IL-1β, pro-IL-18 and GSDMD into their active forms. Little is known of how inflammasome signalling is silenced *in vivo*, and the importance of inflammasome silencing to enforce or restore tissue homeostasis^2^. Using *in vitro* techniques, we discovered how CASP1 activity is switched off in inflammasome-signalling cells. Using ectopically-expressed CASP1 in *Casp1*-deficient macrophages, we discovered that CASP1 activation triggers its timed deactivation through a proteolytic mechanism that we called the ‘CASP1 proteolytic timer’^3^ (**Suppl. Fig. 1A**). CASP1 activation within inflammasomes first involves the recruitment of full-length (p46) CASP1 monomers to the inflammasome, after which these monomers cluster for the proximity-induced dimerisation of the CASP1 protease domain, thereby generating the active protease fold within p46 dimers^3^. CASP1 p46 dimers then self-process at several sites within the protease domain to further activate CASP1 in a species termed p33/p10 dimers^3,44^. CASP1 p33/p10 dimers are bound to the inflammasome and cleave substrates (e.g. pro-IL-1β, pro-IL-18 and GSDMD)^3^. Importantly, the activity of CASP1 p33/p10 dimers is controlled by a second self-cleavage event, in which CASP1 autoprocesses the CDL linker; this removes the CARD domain from the CASP1 protease domain, and consequently releases the p20/p10 dimeric protease domain from the inflammasome. This dimer is inherently unstable when it is not anchored to the inflammasome, and so the dimer dissociates into monomers, thereby terminating CASP1 protease activity^3^. Thus, CASP1 activation within inflammasomes is intrinsically coupled to its deactivation, with proteolytic mechanisms elegantly controlling both of these events in cells.

In the current study, we examined whether this self-limiting ‘proteolytic timer’ silences signalling by endogenous CASP1, and how this may influence *in vivo* tissue functions in homeostasis and organ challenges that can precipitate disease. For this, we generated a homozygous knock-in mouse model harbouring a compound point mutation at the CDL linker of the endogenous *Casp1* locus (E102N/D103N; *Casp1.CDL*) to break the off-switch of the CASP1 proteolytic timer. We confirmed that this mutation suppressed CDL cleavage of endogenous CASP1, and caused heightened CASP1 protease activity and boosted release of mature IL-1β (**Fig. 1, Supp Fig. 1**). Of note, this CDL mutation does not generate a constitutively active CASP1 protease; rather, CASP1 still requires an inflammasome-activating stimulus to acquire protease activity. These *in cellulo* studies with endogenous CASP1 confirmed our previous studies with ectopic CASP1 that CASP1 CDL auto-cleavage prevents inflammasome hyperactivity by terminating CASP1 proteolytic signalling.

Confirming that the CASP1 proteolytic timer silences endogenous inflammasome signalling allowed us to investigate potential functions for this ‘off-switch’ during *in vivo* homeostasis and disease. Inflammasome signalling is implicated as a major driver of anxiety and neurodegeneration in mice and humans^4–6,11,45,46^, and our data support this causal link. We found that under homeostatic conditions, CASP1 CDL mutation to disable the proteolytic timer ‘off switch’ promoted an anxiety-like phenotype in aged *Casp1.CDL* mice (**Fig. 2A-D**, **Supp Fig 3**). Further, inflammasome hyperactivity in *Casp1.CDL* mice enhanced amyloid-induced spatial learning deficits in a murine genetic neurodegenerative model (**Fig. 2E-F**). Inflammasome signalling *in vivo* has been generally studied in response to major tissue challenges, such as the accumulation of misfolded proteins in the brain, or metabolic disturbances. Our observation that signal-induced inflammasome hyperactivity affects brain functions that control anxiety-like behaviour in the absence of an overt challenge suggests that everyday homeostatic fluctuations in tissue biology provoke inflammasome signalling, which must then be silenced to reinforce steady-state tissue homeostasis.

Our studies of steady-state haematopoiesis in *Casp1.CDL* mice also support this notion of inflammasome activation and timely deactivation during everyday homeostatic challenges. Here, CASP1 CDL mutation boosted bone marrow granulopoiesis without affecting monocyte production, leading to greater neutrophil egress into the blood and the spleen (**Fig. 3, Supp Fig. 4**). The enhanced presence of neutrophils in the spleen in steady-state is in keeping with a recent report that neutrophils exit blood vessels to enter tissues via integrin-induced, sub-lytic NLRP3 inflammasome signalling^47^. It is well accepted that hematopoietic stem cells provide a “demand-adapted axis” in the bone marrow to replace mature immune cells during an acute inflammatory response^48,49^; here, IL-1:IL-1R1 in the non-hematopoietic compartment elicits the production of G-CSF, which triggers emergency granulopoiesis^12^. However, in our studies, WT and *Casp1.CDL* mice were not subjected to an overt challenge, were housed in SPF conditions without noticeable indicators of infection, and did not show differences in serum G-CSF. Our data thus suggest that CASP1 is activated and must be deactivated in the steady state, to prevent mild granulopoiesis and neutrophil egress to the circulation and tissues. This did not reflect an intrinsic property of *Casp1.CDL* hematopoietic progenitor cells, because these formed granulocyte progenitors at a similar rate to WT cells in *ex vivo* colony forming unit assays. Altered granulopoiesis in *Casp1.CDL* mice may reflect the paracrine effects of low-level inflammasome cytokines (e.g. IL-1β) on hematopoietic or myeloid progenitor cells^14,15^, or a G-CSF-independent mechanism of demand-induced granulopoiesis caused by increased entry of neutrophils into tissue^47^.

Inflammasome signalling drives chronic low-grade inflammatory responses that contribute to diverse diseases of almost all body systems, including neurodegenerative diseases, cardiovascular diseases, and metabolic dysfunction-associated steatotic liver disease (MASLD). Approximately 20-30% of Western society is currently affected by MASLD, and this proportion is expected to rise steeply in future (25% increase by 2030)^50,51^. Disease progression is generally slow, with people with MASH taking an average of 7 years to progress one fibrosis stage^52,53^. Clinical diagnostics for early disease progression are limited, causing major delays in detecting this slowly progressing disease. Growing evidence implicates chronic inflammation as a major driver of disease progression from hepatic steatosis to liver fibrosis^54,55^. While pathological functions for inflammasomes in driving liver dysfunction and diet-induced peripheral inflammation are well documented^42,56–59^, mechanisms that restrain pathological liver inflammation, and immune resolution mechanisms to promote liver healing are poorly understood. MASLD is characterised by excessive lipid accumulation in the liver, and this is reflected in the MCD diet-induced liver disease model we employed in this study. Mice fed an MCD diet for 2 weeks, regardless of genotype, accumulated lipids such as diacylglycerol (DG), triacylglycerol (TG) and sphingolipids, and this was associated with early indicators of MASLD such as steatosis (**Fig. 4**).

Inflammasome hyperactivity in *Casp1.CDL* mice accelerated MASLD disease progression from steatosis to MASH, with accompanying hepatocyte ballooning, inflammation, and liver damage (**Fig. 4**). Spatial transcriptomics identified a spot cluster (cluster 2) that was more abundant when mice were fed the MCD diet, and further enriched by CASP1 CDL mutation. This spot cluster displayed an innate immune-like-signature, including expression of genes linked to IL-1R signalling and its downstream transcriptional effector, NF-*κ*B. IL-1 is well established to trigger chemokines that direct the recruitment of myeloid cells, such as monocytes. In keeping with this, the MCD diet induced the infiltration of macrophages into the liver. Spatial transcriptomics analyses suggested that infiltrating macrophages are proximal to resident macrophages and hepatocytes in WT MCD-fed mice, while inflammasome hyperactivity uncouples these spatial arrangements and the cell-cell communications they enable in *Casp1.CDL* mice. While it is possible that this finding reflects sample bias, for example introduced by hepatocyte death or the presence of lipid droplets, this is unlikely to be the case as it specifically affected macrophage-hepatocyte relationships and not other cell-cell relationships within the tissue. Given that CASP1 CDL mutation increased diet-induced macrophage infiltration, and hyperactive inflammasome signalling in macrophages would be expected to boost IL-1 release, is it possible that the heightened inflammatory milieu in CASP1 CDL mice sensitised hepatocytes to lipid-induced cell death^37^, thereby uncoupling hepatocytes from their surrounding macrophages.

The remarkable regenerative capacity of the liver allowed us to investigate how CASP1 inflammatory programs are silenced to restore homeostasis after the removal of a major challenge to organ function. After MCD feeding for 4 weeks to induce advanced liver inflammation and damage, we switched mice to a healthy diet and monitored liver healing (**Fig. 7**). CASP1 CDL mutation prolonged the presence of hepatic infiltrating macrophages as well as the clinical indicators of liver damage and dysfunction. CASP1 CDL self-cleavage to switch off CASP1 signalling is thus critical to silence hepatic inflammatory responses and restore normal liver function.

Current MASLD management strategies solely rely on lifestyle interventions (e.g. healthy diet, exercise) aiming to reduce body mass. The clinical management of MASLD rarely involves anti-inflammatory therapies or monitors the inflammatory status of people living with MASLD. Our data reveal the importance of CASP1 inactivation for suppressing hepatic inflammation and promoting liver healing. With inflammasome inhibitors rapidly advancing through clinical trials as new anti-inflammatory agents, we propose that inflammatory status should be a clinical consideration for treating MASLD, including with upcoming new inflammasome-targeting therapeutics.

In sum, our data provide the first *in vivo* mechanism for inflammasome signal termination. Here, the intrinsic proteolytic timer function of CASP1 self-limits inflammasome activity to enforce homeostasis in the steady state, temper inflammatory programs during organ challenge, and reinstate tissue homeostasis following the removal of a major challenge to organ function. Inflammasome signal termination by CASP1 CDL cleavage thereby ensures the timely resolution of tissue inflammation. The critical importance of CASP1 silencing for homeostasis and restoring tissue function highlights CASP1 as an emerging target for new anti-inflammatory drugs.

## Resource availability

### Lead contact

Further information and requests for resources and reagents should be directed to and will be fulfilled by Dr. Sabrina Sofia Burgener and Prof. Kate Schroder

### Materials availability

Materials and reagents generated in this study are available upon a reasonable request from the lead contacts and may require a completed Material Transfer Agreement.

### Data and code availability

- All data reported in this paper will be shared by the lead contacts upon request
- Codes for analysing spatial transcriptomics are available at https://github.com/jipsi/spatial_casp. Raw FASTQ and spatial files will be provided under a GSE accession number upon manuscript acceptance.
- Any additional information required to reanalyse the data reported in this study are available from the lead contacts upon request

## Acknowledgments

This work was supported by the MRFF Project Breaking Through Dementia (MRRF GA39196 to JG), the Clem Jones Centre for Ageing Dementia Research (to JG and KS), and the National Health and Medical Research Council of Australia (NHMRC) (1194329 to AJM, 2009075 to KS). SSB was supported by a Swiss National Science Foundation Postdoc Mobility Fellowship (P2BEP3_191800) and a Novartis Foundation for Medical-Biological Research Fellowship (21C133). SSB and UB were awarded a 10x Genomics-Illumina Visium Spatial Biology Kickstarter Award. MTM was supported by the Yulgilbar Alzheimer’s Foundation PhD stipend. QN and LGB are supported by the NHMRC (2008928 to QN, 2030460 to LGB). DB was supported by the MRC (MR/Z504221/1) and BBSRC (BB/Y009703/1). SD and PMK were supported by a Wellcome Investigator Award (224290/Z/21/Z to PMK). The authors acknowledge the facilities, and the scientific and technical assistance, of Microscopy Australia at the Centre for Microscopy and Microanalysis at The University of Queensland. All behavioural work was conducted at the Queensland Brain Institute Animal Behaviour Facility. The authors thank Onkar Mulay, Jared R Coombs, Tyron Esposito and Drs Caroline L Holley, Grace MEP Lawrence and Stefan Emming for technical assistance, and Dr Madhavi Maddugoda for critical reading of early manuscript drafts.

## Author contributions

SSB and KS conceived the study and acquired project funding. SSB designed and performed experiments, analysed data, initiated and managed collaborations (SD, PM, GM, AC, AM, PMK), managed mouse ethics and colonies, prepared all figures for the submission and wrote the manuscript draft. MTM performed and analysed behaviour studies, with input from DGB, LGB and JG. SD analysed spatial-transcriptomics experiments with input from SSB and PMK. PKM performed and analysed lipidomic data with input from SSB and TJCC, PM and AJM. EF, KMK, MDO gave technical support and advice. UB, ZX and QN performed the spatial transcriptomic run with input from SSB. GM and AC performed blind scoring of liver H&E section provided by SSB. DB gave technical support and advice. KS supervised the project and wrote the final manuscript with SSB. All authors discussed experimental results and gave input into the manuscript.

## Materials and Methods

### Key resource table

| REAGENT or RESOURCE | SOURCE | IDENTIFIER |
| --- | --- | --- |
| **Antibodies** |  |  |
| CD45 FITC | BioLegend | Clone 30-F11,<br>Cat.#103108;<br>RRID: AB_312973 |
| CD45 BV510 | BioLegend | Clone: 30-F11,<br>Cat.# 103138; RRID:<br>AB_2563061 |
| CD11b BV421 | BioLegend | Clone: M1/70,<br>Cat.#101236;<br>RRID: AB_11203704 |
| CD11b APC | BioLegend | Clone: M1/70,<br>Cat.#101212;<br>RRID: AB_312795 |
| CD11c APC-Cy7 | BioLegend | Clone: N418,<br>Cat.#117323;<br>RRID: AB_830646 |
| CD68 BV605 | BioLegend | Clone: FA-11,<br>Cat.#137021;<br>RRID: AB_2616811 |
| I-A/I-E (MHCII) APC | BioLegend | Clone:M5/114.15.2,<br>Cat.#107614;<br>RRID:AB_313329 |
| CD115 BV711 | BioLegend | Clone: AFS98<br>Cat.#135515<br>RRID: AB_2562679 |
| CD115 APC | BioLegend | Clone: AFS98<br>Cat.#135510<br>RRID: AB_2085221 |
| Ly6C BV785 | BioLegend | Clone: HK1.4<br>Cat.#128041<br>RRID: AB_2565851 |
| Ly6C PE | BioLegend | Clone: HK1.4<br>Cat.#128007<br>RRID: AB_1186133 |
| Ly6G PE-Cy7 | BioLegend | Clone: 1A8<br>Cat.#127618<br>RRID: AB_1877261 |
| Sca-1 PE-Cy7 | BioLegend | Clone: D7<br>Cat.#1008114<br>RRID: AB_493596 |
| CD16/32 BV421 | BioLegend | Clone: 93<br>Cat.#101331<br>RRID: AB_2562188 |
| Gr-1 (Ly6C/Ly6G) APC | BioLegend | Clone: 6B6-8C5<br>Cat.#108412<br>RRID: AB_313377 |
| CD45R (B220) FITC | BioLegend | Clone: RA3-6B2<br>Cat.#103206<br>RRID: AB_312991 |
| CD45R (B220) APC | BioLegend | Clone: RA3-6B2<br>Cat.#103212<br>RRID: AB_312997 |
| CD44 BUV395 | BD Bioscience | Clone: IM7<br>Cat.#740215<br>RRID: AB_2739963 |
| CD62L PE-Cy5 | BioLegend | Clone: MEL-14<br>Cat.#104410<br>RRID: AB_313097 |
| Ter119 APC | BioLegend | Clone: TER-119<br>Cat.#116212<br>RRID: AB_313713 |
| CD3e APC | BioLegend | Clone: 145-2C11<br>Cat.#100312<br>RRID: AB_312677 |
| CD117 APC-Cy7 | BioLegend | Clone: 2B8<br>Cat.#105825<br>RRID: AB_1626280 |
| CD135 PE | BioLegend | Clone: A2F10<br>Cat.#135306<br>RRID: AB_1877217 |
| CD34 FITC | ThermoFisher (eBioscience) | Clone: HM34<br>Cat.#11-0341-85<br>RRID: AB_465022 |
| FVS700 | BD Bioscience (Becton, Dickinson & Company) | Cat.#564997;<br>RRID: AB_286937 |
| CD16/32 (clone 2.4G2) | BD Bioscience (Becton Dickinson & Company) | Clone: 2.4G2,<br>Cat.#553142;<br>RRID: AB_394657 |
| CD68 | Abcam | Cat.#ab125212<br>RRID: AB_10975465 |
| Clec4f | R&D Systems | Cat.#AF2784<br>RRID: AB_2081339 |
| Ly6G | BioXCell | Clone: 1A8<br>Cat.#BE0075-1<br>RRID: AB_1107721 |
| αSMA | Cell Signaling Technology | Cat.#19245<br>RRID: AB_2734735 |
| ASC | Cell Signaling Technology | Cat.#67824<br>RRID: AB_2799736 |
| CASP1 (Casper-1) | Adipogen | Cat.#AG-20B-0042-C100<br>RRID: AB_2490248 |
| IL-1β | R&D Systems | Cat.#AF-401-NA<br>RRID: AB_416684 |
| GSDMD | Abcam | Cat.#ab209845<br>RRID: AB_2783550 |
| SYTOX green nucleic acid stain | Thermo Fisher (Invitrogen) | Cat.#S7020 |
| Propidium Iodide | Biolegend | Cat.#421301 |
| DAPI (4',6-diamindino-2-phenylindole) | Sigma | Cat.#D9542-5MG |
| Alexa Fluor 647 Phalloidin | Thermo Fisher (Invitrogen) | Cat.#A22287;<br>RRID: AB_2620155 |
| Goat anti-Rabbit IgG (H+L) Cross-Adsorbed Secondary Antibody, Alexa Fluor Plus 594 | Thermo Fisher (Invitrogen) | Cat.#A32740;<br>RRID: AB_2762824 |
| Chicken anti-Goat IgG (H+L) Cross-Adsorbed Secondary Antibody, Alexa Fluor 488 | Thermo Fisher (Invitrogen) | Cat.#A21467;<br>RRID: AB_2535870 |
| Goat anti-Rat IgG (H+L) Cross-Adsorbed Secondary Antibody, Alexa Fluor 488 | Thermo Fisher (Invitrogen) | Cat.#A11006;<br>RRID: AB_2535870 |
| Donkey anti-goat IgG-HRP linked | Jackson ImmunoResearch | Cat.#705-035-003<br>RRID: AB_2340390 |
| Horse anti-mouse IgG-HRP linked | Cell Signaling Technology | Cat.#7076<br>RRID: AB_330924 |
| Goat anti-rabbit IgG-HRP linked | Cell Signaling Technology | Cat.#7074S<br>RRID: AB_2099233 |
| **Experimental models: organisms/strains** |  |  |
| C57BL/6J | Ozgene ARC – Jackson Lab | RRID: IMSR_JAX:000664 |
| *Casp1.CDL* | Generated in-house using CRISPR, on C57BL/6J background | N/A |
| **Chemicals, peptides, and recombinant proteins** |  |  |
| Macrophage-colony stimulating factor (M-CSF), endotoxin-free | Generated by the UQ Protein Expression Facility in insect cells, confirmed to be endotoxin free (alternative commercial supplier: Peprotech) | N/A<br>(Cat.# 315-02) |
| PBS (Ca<sup>2+</sup>Mg<sup>2+</sup> free) | Gibco | #10010023 |
| EDTA (0.5M) | Gibco | #15575020 |
| Trypan blue | Sigma | #T8154-100ML |
| DMEM | Thermo Fisher | #11995073 |
| RPMI 1640 | ThermoFisher | #11875085 |
| Sodium pyruvate | ThermoFisher | #11360070 |
| Foetal bovine serum (FBS) | ThermoFisher | #10099141 |
| Penicillin/streptomycin | Gibco | #15140122 |
| GlutaMAX | Gibco | #35050061 |
| 0.25% Trysin-EDTA | Thermo Fisher | #25200072 |
| Bovine serum albumin (BSA) fraction V (7.5%) | Thermo Fisher | #15260037 |
| FixPerm buffers | BioLegend | #426803 |
| Pierce™ 16% Formaldehyde (w/v), Methanol-free | Thermo Fisher | #28908 |
| ProLong™ Gold Antifade Mountant | Thermo Fisher | #P36934 |
| Bovine serum albumin (BSA) | Scientifix | #BSASAU |
| Digitonin | Sigma-Aldrich | #D141-100MG |
| DL-Dithiothreitol | Sigma-Aldrich | #D9779 |
| DPX mountant for histology | Merck Millipore | #06522 |
| Fluorescence mounting medium | Agilent | #S302380-2 |
| Nonidet P 40 Substitute (NP-40) | Sigma-Aldrich | #74385 |
| NuPAGE™ LDS Sample Buffer (4 X) | ThermoFisher | #NP0008 |
| PhosSTOP™ | Merck Millipore | #4906845001 |
| cOmplete™, Mini, EDTA-free Protease Inhibitor Cocktail | Merck Millipore | #11836170001 |
| Clarity Western ECL Substrate | BioRad | #1705061 |
| SuperSignal West Femto Maximum Sensitivity Substrate | ThermoFisher | #34095 |
| ATX Ponceau S red staining solution | Merck Millipore | #09189-6X1L-F |
| Stripping Buffer |  |  |
| 5,5'-Tetramethylbenzidine (TMB) | Merck Millipore | #CL07 |
| Ultrapure *Escherichia coli* K12 LPS | Invivogen | #tlrl-eklps |
| Nigericin from *Streptomyces hygroscopicus* | Sigma-Aldrich | #N7143 |
| ATP | Sigma-Aldrich | #A2383 |
| PA-1 | Quench (alternative commercial supplier: Invivogen) | #tlrl-rod |
| LFn-Fla | Quench (alternative commercial supplier: Invivogen) | #tlrl-rod |
| Biotin-VAD-FMK | MedChemExpress | #HY-100894 |
| Ac-WEHDI-AFC | SantaCruz | #sc-311282 |
| **Critical commercial assays** |  |  |
| Methionine-Choline-supplemented (MCS) diet | Speciality Feed | #SF13-137 |
| Methionine-Choline-deficient (MCD) diet | Speciality Feed | #SF13-136 |
| MethoCult | StemCell Technologies | #M3434 |
| Click-iT EdU Kit for Flow Cytometry | ThermoFisher | #C10425 |
| CytoTox 96 non-radioactive Cytotoxicity Assay LDH | Promega | #G1780 |
| IL-1 beta/IL1F2 DuoSet ELISA | R&D Systems | #RDSDY401 |
| BCA Total Protein | Pierce Thermo Fisher | #23225 |
| 4-20% Mini-PROTEAN TGX Stain-Free Protein Gels | BioRad | #4568094 |
| Paramagnetic streptavidin beads | Promega | #Z5481 |
| **Software and algorithms** |  |  |
| FlowJo | BD (Becton, Dickinson & Company) | https://www.flowjo.com/ |
| GraphPad Prism | GraphPad | https://www.graphpad.com/scientific-software/prism/ |
| Fiji | ImageJ (version 2.00-rc 69/1.52p) | https://imagej.net/software/fiji/ |
| spaceranger | https://www.10xgenomics.com/support/cn/software/spaceranger/latest | v1.3.0 |
| Cell2location | https://github.com/BayraktarLab/cell2location | v0.1 |
| Seurat | https://satijalab.org/seurat/ | v4.3.0 |
| R | https://www.r-project.org/ | v4.2.2 |
| Python | https://www.python.org/ | v3.8 |
| **Other** |  |  |
| T25 cell culture grade flasks | Thermo Fisher | # 156340 |
| 70µm cell strainers | Bio-Strategy | #BDAA352350 |
| FACS tubes (5ml round bottom polystyrene test tube) | In Vitro Technologies | #352052 |
| Cell scraper | Sarstedt | #83.3951 |
| 12 mm coverslips | Bio-Strategy | #EPBRCSC121GP |
| Microscope slides | Bio-Strategy | #EPBRSF21201 |
| 24-well tissue culture plates | Sigma-Aldrich | #3526 |
| Biosafety cabinet | Gelaire | #BHEN2004D |

### Data and code availability

- All data reported in this paper will be shared by the lead contacts upon request
- Code used for the analysis of spatial transcriptomics data is available at https://github.com/jipsi/spatial_casp. Raw FASTQ and spatial files will be provided under a GSE accession number upon manuscript acceptance for publication.
- Any additional information required to reanalyse the data reported in this paper is available from the lead contacts upon requests

### Animals

All animal studies were approved by the University of Queensland (UQ) Animal Ethics Committee and conducted following federal legislation on animal welfare (2023/AE000019, 2023/AE000020, 2022/AE000351, 2019/AE000280, 2019/AE000281, 2023/AE000186). Mice were housed in specific pathogen-free (SPF) facilities in individually ventilated and autoclaved cages with 12/12-hour light/dark cycles, autoclaved acidified water, food, bedding and environmental enrichment. Each set of experimental data indicates mouse age and sex in the figure legend. C57BL/6J breeders were originally sourced from Ozgene or the Australian Resource Centre, and bred in-house, including for the generation of *Casp1.CDL* mice (see below). *APP23* mice express the human APP751 Swedish double mutation (KM670/671NL) under the control of the neuron-specific mThy1.2 promoter^7^. *APP23* mice were crossed with *Casp1.CDL* mice to generate *App23.Casp1.CDL* mice, and all comparisons between these mice were performed with littermates.

### Generation of *Casp1.CDL* mice using CRISPR

*Casp1.CDL* (E102N/D103N) mice were generated by microinjection of wild-type zygotes with ribonucleoprotein (RNP) complexes^60^ by the Transgenic Animal Service of Queensland (TASQ)-QFAGE CRISPR service at the University of Queensland. Recombinant Cas9 nuclease, sgRNA and nuclease-free duplex buffer were purchased from IDT. HDR donor template of the genomic sequence of CASP1 on Chr 9 encompassing sequence ranging from exon 1 to exon 7 including the E102N/D103N compound mutation, together with sgRNA sequence CACCGTTCTAAAGGGCAAAACTTGA were used (**Suppl. Fig 1B**). Microinjection into the male pronucleus of 185 zygotes was performed, and 135 live 2-cell stage embryos were transferred into the infundibulum of pseudopregnant CD1 females using standard protocols. 15 founders were screened by PCR and T7 endonuclease assay, and at least 5 mice had the correct desired mutation. Founders were crossed with C57BL/6J mice and mutated alleles in F1 mice were identified by DNA sequencing (**Supp. Fig. 1C**). F1 mice with the E121N/D122N mutation were intercrossed to generate homozygote *Casp1.CDL* mice. F2 progeny and their littermates were used in this study. Genotyping was performed using primers F1 5’CACATTGGCCTCAGTCTCAC’3 and R1 5’ACACAAGAACTGCCTTGCTC’3 resulting in a 295bp PCR product. PCR cleanup (ExoSAPit) was performed and sequenced the PCR product using BigDye Terminator v3.1 Sanger sequencing, using a sequencing cleanup by Agencourt CleanSeq magnetic beads, and analysed the sequencing reaction products on a 3730xl DNA analyser (**Suppl. Fig 1B-C**).

### Behaviour studies

For all behavior experiments, male and female mice were separated and aged to 18 months or less. For experiments using *App23* and *App23.Casp1.CDL* mice, mice were aged to 12 months. Mice were habituated to the researcher for at least 5 days before commencing behavioral testing. For all tests, mice were habituated in the room in which the test was performed for at least 30 minutes before testing. The weight of all animals was monitored weekly.

### Open-field test

Open-field test was performed to evaluate general mouse behavior. Each mouse was placed inside a well-lit, square-shaped box (60 to 70 lx) in which animal movement, position and speed were monitored by infrared beam breaks that project across the open field along the X, Y, and Z axes. Mice were placed in the open field box and allowed to explore for 30 min. The software was set up according to the General Open Field test from the Activity Monitor version 7 manual (MED Associates, Inc.). All data was analyzed using Activity Monitor 7 software, SOF-812 (MED Associates, Inc.).

### The elevated plus maze (EPM)

The EPM was used to evaluate anxiety-like behavior in mice. Briefly, mice were placed in the central area of the maze (elevated cross-shaped apparatus with a central square, closed arms, and open arms with unprotected edges). Mice were allowed to explore the maze for 5 min (80-100 lx) and their movements were recorded using an overhead camera with EthovisionXT^TM^ tracking software (Ethovision). Videos were analyzed using EthovisionXT^TM^ software to calculate percentage of time spent in each EPM arm.

### The active place avoidance (APA)

The APA test was used to assess hippocampal-dependent spatial learning and memory^61^. Briefly, mouse behaviour was assessed over 6 days in an elevated, rotating circular arena with a transparent, 32 cm high perimeter circular fence (BioSignal Group) with a metal grid floor, enclosing a total diameter of 77 cm. Four high-contrast visual cues were placed at 90° from each other around the arena. The arena and floor were rotated at a speed of 1 rpm, with a mild shock (500 ms, 60 Hz, 0.5 mA) delivered through the grid floor each time the mouse entered the 60-degree shock, then every 1500 ms until the animal left the shock zone. The shock zone was maintained at a constant position relative to the room and had one associated visual cue. Arena rotation was designated clockwise or anti-clockwise, and the room lighting was set to 30 lx.

On day 0, mice were habituated in the arena for 5 min with the shock turned off and rotation turned on. From day 1 to day 5, the shock and rotation were both turned on. The test was performed for 10 min per day over five consecutive days. Recorded tracks were analyzed with Track Analysis software (BioSignal Group). The number of shocks, number of entries to the shock zone, time to first entry, time to second entry, and maximum avoidance time for each mouse were compared over the days of testing. To standardize results, performance at the start of the test phase to the end of the test phase was compared by calculating: (d5 value – d1 value)/d1 value x 100. This was calculated for shock number and maximum avoidance time, then averaged across the 5 parameters to obtain “Cognitive Index”.

### *In vivo* model of MASLD

MASLD is a sexually dimorphic disease with more robust phenotypes in male rodents, mirroring its greater prevalence in men versus pre-menopausal women^62^. C57BL/6J and *Casp1.CDL* male mice, of 12 weeks of age, were randomly divided into groups (n=2-3 mice per group, n=4-5 repetition) and fed a methionine-choline-supplemented (MCS) diet (Speciality Feed, #SF13-137) or methionine-choline deficient (MCD) diet (Speciality Feed, #SF13-136) for 2 or 4 weeks. Disease resolution was assessed by feeding a methionine-choline deficient (MCD) diet for 4 weeks, followed by 7 days of an MCS diet. Mouse body weights were recorded twice a week for all cohorts.

### Liver histology

Mouse livers were isolated and cut in random 3-4 mm pieces. For each genotype or treatment group, three random liver pieces per mouse were selected, fixed with 4% PFA and paraffin-embedded (Translational Research Institute Histology Facility). Sections were sliced (thickness 5µm) and mounted on SuperFrost Plus slides. The livers were blindly assessed for steatosis, hepatic ballooning, lobular inflammation, fibrosis, and NAS scores by pathologists from Envoi Pathology, Dr Greg Miller and Prof. Andrew Clouston.

### Liver immunohistochemistry

Immunohistochemistry was performed to stain liver sections for macrophages (anti-CD68, 1:400, Abcam), and neutrophils (anti-Ly6G, 1:400, BioXCell). Slides were dewaxed by baking for 30 min at 60°C followed by cooling for 5 min at RT. Antigen retrieval was performed using citrate buffer at full power in a microwave until boiling, followed by low power (10%) for 15 min, which was blocked in histology blocking buffer for 1 h at RT. Sections were incubated with primary antibody overnight at 4°C. Intrinsic peroxidase activity was removed using 3% H_2_O_2_ in TBS + 0.05% Triton-X. The secondary mouse IgG biotinylated antibody was added for 1.5 h at RT. ABC reagent was added for 30 min at RT, and DAB substrate was added to slides and monitored for development by light microscopy. Slides were washed once with MilliQ water for 5 min and counterstained with haematoxylin for 1 min. Slides were dehydrated in 70% ethanol, 95% ethanol, 100% ethanol, and xylene and mounted in DPX.

### Hematology analysis

Mice were bled retro-orbitally, and blood was collected in an EDTA tube. A Mindray Hematology Analyser quantified WBC, RBC, Haemoglobin, Haematocrit and platelets in whole blood.

### Serology liver enzymes and FIB-4 score

Serum was collected from full blood, and analysed by the UQ Clinical Pathology School of Veterinary Science to assess levels of albumin, cholesterol, triglycerides, urea, total bilirubin, creatinine, alkaline phosphatase (ALP) and alanine aminotransferase (ALT), and aspartate aminotransferase (AST). FIB-4 score was calculated using the following formula for all animals, using 55 years of age to approximate murine age to human years:

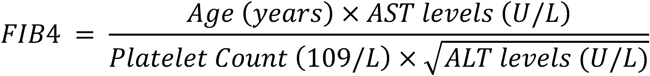

### Liver tissue homogenisation

Liver tissues (20–40 mg wet weight) were homogenized in 200 µl PBS (without calcium or magnesium) with a tissue lyser (Qiagen). Extracts were then sonicated (25 s at 13% amplitude; Misonix S-4000 Sonica). Protein concentrations of the tissue samples were determined using Qubit Protein assay kit (#Q33212), and 50 μg of each sample was aliquoted. To prepare for lipid extraction, the tissue homogenates were stored at -80°C for 2 hours and dried overnight using Savant SpeedVac (Thermofisher).

### Lipid extraction of liver homogenates

Samples were randomised and single-phase butanol/methanol extracted lipids. As previously described, 100 µl of butanol:methanol containing internal standard mixtures was added to each sample. Samples were vortexed, sonicated in a water bath sonicator for an hour and centrifuged for 10 min at 13,000*g* to precipitate proteins from lipid extracts. The supernatant was transferred into 0.2 ml micro-inserts in glass vials to be stored at -80°C until mass spectrometry analysis. In addition to sample extract and PQC, technical QC samples (TQC) consisting of PQC extracts which were pooled before dividing into individual vials, were added to the run (at a ratio of 1 TQC per 20 samples) to provide a measure of technical variation derived from the mass spectrometer. Changes in peak area, width and retention time were monitored between TQCs to determine the performance of the LC-MS/MS analysis and were subsequently used to account for any differential responses across the analytical batches. The structural elucidation of some phospholipid lipid species was acquired by running plasma samples and/or synthetic standards under the same chromatographic conditions and running orthogonal mass spectrometry experiments (negative mode MS/MS, lithium adduct MS/MS)^63^. Lipid species were relatively quantified by comparison to the relevant internal standard.

### Liquid chromatography MS/MS

Lipid analyses were performed as described previously^63^ with modifications. Lipid extracts were analysed on an Agilent 6495C QQQ mass spectrometer with an Agilent 1290 series HPLC system and a single ZORBAX eclipse plus C18 column (2.1×100mm 1.8mm, Agilent) with the thermostat set at 45°C. Dynamic scheduled multiple reaction monitoring in positive and negative ion mode measured a total of 665 lipid species (Supplementary Table 1). The running solvent consisted of solvents A and B containing of water: acetonitrile:isopropanol in the ratios of 5:3:2 and 1:9:90 respectively, including 10 mM ammonium formate (in solvents A and B) and 5 μM medronic acid (in solvent A only). The chromatography and mass spectrometer settings are shown in Supplementary Tables 1 and 2. The isolation widths for Q1 and Q3 were set to “unit” resolution (0.7 amu).

**Supplementary Table 1.** Chromatographic settings and conditions on the Agilent 1290 Infinity/Infinity II LC.

|  |  |  |  |
| --- | --- | --- | --- |
| Analytical column | Agilent Zorbax Eclipse Plus C18, 1.8 µm, 100 x 2.1 mm column, p/n 959758-902 |  |  |
| Inline filter | Agilent 1290 Infinity II inline filter, 0.3µm, p/n 5067-6189 |  |  |
| Column temperature | 45 °C |  |  |
| Injection volume | 1 µL |  |  |
| Autosampler temperature | 20 °C |  |  |
| Mobile phase | Solvent A: 50% water, 30% acetonitrile, 20% 2-propanol, 10 mM ammonium formate, 5 µM medronic acid<br>Solvent B: 90% 2-propanol, 9% acetonitrile, 1% water, 10 mM ammonium formate |  |  |
| Flow rate | 0.4 mL/min |  |  |
| Gradient | Time (min) | % A | % B |
|  | 0 | 85 | 15 |
|  | 2.5 | 50 | 50 |
|  | 2.6 | 43 | 57 |
|  | 9 | 30 | 70 |
|  | 9.1 | 7 | 93 |
|  | 11 | 4 | 96 |
|  | 11.1 | 0 | 100 |
|  | 12 | 0 | 100 |
|  | 12.2 | 85 | 15 |
|  | 16 | 85 | 15 |

**Supplementary Table 2.** Instrument parameters for Agilent 6495C triple quadrupole LC/MS.

|  |  |
| --- | --- |
| Parameter | Agilent 6495C |
| Ion source | Agilent Jet Stream |
| Polarity | Positive and negative switching |
| Gas temperature | 150 °C |
| Drying gas (nitrogen) | 17 L/min |
| Nebuliser gas | 20 psi |
| Sheath gas | 200 °C |
| Sheath gas flow | 10 L/min |
| Capillary voltage | 3,500 V (+), -3,000 V (-) |
| Nozzle voltage | 1,000 V (+), -1,500 V (-) |
| iFunnel high/low pressure RF | 200/110 V (+), 150/60 V (-) |
| Scan type | Dynamic MRM (DMRM) |
| Q1/Q2 resolution | Unit (0.7 amu) |
| Delta EMV | 100 V (+), 100 V (-) |
| Cell acceleration voltage | 4-5 V |
| Cycle time | 650 ms |
| Time filter width | 0.07 min |
| Total number of MRMs | 665 (Positive: 646, negative: 19) |
| Fragmentor | 166 |

### Lipid nomenclature

The lipid names used in this study follow guidelines set by the LIPID MAPS consortium^64–67^. Phospholipids with detailed characteristics (that is, acyl chain composition) are annotated as ‘PC(16:0_20:4)’, with PC being the lipid class and (16:0_20:4) representing the acyl chains found on the glycerol backbone, irrespective of sn1 or sn2 position. Lipids without specific structural annotations are named based on their sum acyl chain length and degrees of saturation (for example, PC(36:4)).

### Lipid data normalization

Analyte areas were obtained from integrating chromatograms that corresponded to a lipid of interest using the MassHunter Quantitative Analysis Software (version B9.00; Agilent). In brief, individual analyte areas were divided by the area of the corresponding ITSDs, and the median of ITSD mixtures containing blank samples was subtracted from each analyte (background subtraction). This value was then multiplied by the ITSD concentration and the individual analyte’s response factor (*R*). Any values that were zeroed after background subtraction as a consequence of being less than the median value of all blank + ITSD samples were replaced with 1/10th of the minimum value for the corresponding analyte. The data were ultimately normalized to pmol or µmol of total lipidome, where the background subtracted datum for an individual lipid was divided by the sum of the total lipidome and multiplied by a factor of 10^6^ for ease of graphical representation.

### Lipidomic analyses

All downstream analyses were conducted in R version 3.6.2 (Comprehensive R Archive Network). UMAP was performed using the FactoMineR and factoextra packages using log10-transformed lipid concentrations. One-way analyses of variance (ANOVAs) followed by post-hoc testing using Tukey’s test were performed using in-built functions in R. ANOVA-generated p values were corrected for multiple comparisons using the Benjamini–Hochberg method for false discovery rate (FDR) (5%) correction. FDR-corrected p values were considered statistically significant when p < 0.05. Z scores were calculated by subtracting the group mean from the individual raw score and dividing by the standard deviation.

### Visium whole spatial transcriptomics tissue processing

Spatial transcriptomics analysis was performed on mouse formalin-fixed paraffin-embedded (FFPE) tissue sections using the 10x Genomics Visium platform. The experimental design comprised twelve samples representing two genotypes and two diet conditions, with one representative liver section with mild (less than 20% steatosis), moderate (20-50% steatosis) and severe (above 50% steatosis) pathology from MCD-fed WT and *Casp1.CDL* mice, alongside three liver sections from MCS-fed controls of each genotype. All samples were multiplexed onto a single Visium slide, with 5μm sections positioned within the designated capture areas marked by fiducial markers.

The slide was processed according to the Visium Spatial Gene Expression Reagent Kit for FFPE protocol (version 1, 10x Genomics). The procedure included hematoxylin and eosin (H&E) staining of the tissue sections, followed by microscopic imaging. After the tissue was de-crosslinked, mouse-specific probes were applied and allowed to hybridize overnight. Following probe extension and release, sequencing libraries were prepared according to the manufacturer’s guidelines. These libraries were then sequenced using the NovaSeq 6000 platform. Associated image files were aligned onto slide-specific fiducials using Loupe browser software (10x Genomics).

### Visium data processing and integration

The resulting FASTQ files were aligned to the mouse genome GRC (mm10) using spaceranger (v1.3.0) software. Loupe browser software was used to align the H&E images with slide-specific fiducials. Each tissue section was annotated, generating tissue x-y coordinate JSON files that were subsequently used by Space Ranger’s ‘count’ function to match gene expression counts with their spatial locations on the tissue.

Seurat (version 4.3.0) was used for data normalization and integration. SCTransform was applied for normalization and scaling, using nCount_Spatial and nFeature_Spatial to regress the counts. All 12 samples were integrated by selecting features, identifying anchors, and finally integrating the data. Cluster memberships were determined using the first 12 principal components at a resolution of 0.3. Before differential gene expression analysis, sequencing depth was accounted for by using PrepSCTFindMarkers() to reverse individual SCT models to use the minimum of median UMI as a covariate for sequencing depth. Wilcoxon rank-sum test was then employed to identify differentially expressed genes, applying a Bonferroni-corrected P-value threshold of 0.05. This analysis was structured to find expression differences between genotypes within each diet group, as well as diet-induced changes within each genotype. Additional pathway analyses were conducted using g:Profiler to infer biological significance of gene lists.

### Cell type deconvolution of Visium spots

Cell type deconvolution within the spatial data was performed using cell2location (v0.1) with single-cell non-parenchymal dataset from an MCD-diet fed mouse model (GSE129516; ^43^). Reference cell type gene expression was modelled through negative binomial regression over 1,000 epochs, followed by mapping spatial gene expression to cellular abundances over 30,000 epochs. Parameter setting included setting N_cells_per_location to 30 and detection_alpha to 20. The predicted cell type abundances, derived from the 5% quantile values of the posterior distribution, were incorporated into the Seurat object as metadata. Cell type abundances were then plotted as bar plots or spatially as a heatmap.

### Bone marrow-derived macrophage (BMDM) preparation

Wildtype and *Casp1.CDL* female and male mice were sacrificed using CO_2,_ and the femurs and tibias were removed. The bones were flushed with sterile PBS + 10% FBS using a 27G needle and 10-ml syringe, then filtered through a 40 µm strainer. The filtered bone marrow was centrifuged at 500 x g for 5 min. The bone marrow pellet was resuspended in 1 ml of RPMI media + 10% FBS + 1% Pen/Strep + 1% Glutamax (BMDM media) per leg (one femur and one tibia) and the volume was adjusted to 60 ml per leg and M-CSF added to a final concentration of 150 ng/ml. 15 ml of the bone marrow suspension was plated into 10-cm sterilin dishes and cultured for five days at 37°C/5% CO_2_. On day 5 in culture, 3 ml of fresh BMDM media containing M-CSF (150 ng/ml) was added to the cells. On day 6, the media was removed from the cells and replaced with ice-cold PBS. BMDMs were harvested from the sterilin dish with an 18G needle and 10-ml syringe. Cells were pelleted and resuspended in BMDM media containing M-CSF (150 ng/ml) for seeding in 96-well plates at a density of 1 x 10^5^ cells/well. Inflammasome assays were performed on day 7.

### Inflammasome assay *in vitro*

To measure NLRP3 or NLRC4 inflammasome signalling *in vitro*, BMDMs were isolated and seeded at a density of 1 x 10^5^ cells/well in a 96-well plate. For NLRP3 inflammasome activation, cells were primed with 100 ng/ml ultrapure *Escherichia coli* K12 LPS in Opti-MEM containing 150 ng/ml CSF for 4 h, followed by nigericin (5 μM) or 2.5 mM ATP for 10 min up to 6 hours. The NAIP-NLRC4 inflammasome was activated using the PA-FlaTox system, which delivers *Legionella pneumophila* flagellin (Fla1) to the cytosol by N-terminal fusion to the *Bacillus anthracis* lethal factor (LFn), and transmembrane transport through the anthrax protective antigen (PA) channel^68,69^. Cells were primed with 100 ng/ml ultrapure *Escherichia coli* K12 LPS in Opti-MEM containing 150 ng/ml CSF for 4 h, followed by 25 ng/mL PA and 0.125 ng/mL Fla1-LFn in Opti-MEM for 10 min up to 6 hours. The supernatant was collected and used for cell death and cytokine assays, and cells were lysed in immunoblotting lysis buffer for protein expression analysis.

### Caspase-1 activity assay *in vitro*

Inflammasome activation was performed as described above. Cell supernatant was replaced with pre-heated 1x Caspase-1 Activity Buffer (Digitonin, DTT, af-WEHD-AFC), and the kinetics of af-WEHD-AFC cleavage by CASP1 was measured at excitation 400 nm and emission 505 nm at 37°C for 60 min.

### Active caspase pulldown

Biotin-VAD-fmk (b-VAD-fmk) was applied to cells to label and trap active CASP1 during inflammasome signalling^3^. BMDMs were primed and stimulated as described. 10µM bVAD-fmk was applied to cells 0.5 h before the inflammasome agonist. bVAD-fmk pull-down was performed on the entire cell output (cell lysate plus supernatant); here, cell supernatant was collected, cells were lysed in bVAD lysis buffer (150mM NaCl, 50mM Hepes pH 8.0, 1% IPEGAL, Protease Inhibitors, Benzonase) and incubated for 5 min on ice and then mixed with cell supernatant. Paramagnetic streptavidin beads were washed and resuspended in PBS + 1% BSA and added to the cell lysate/supernatant mix, and incubated with gentle rotation at 4°C overnight. The streptavidin-unbound fraction was methanol/chloroform-precipitated, and beads were washed three times with buffers with increasing salt concentration (150 – 500mM NaCl, 50mM Hepes pH 8.0, 1% IPEGAL). Beads were then pelleted and resuspended in NuPage LDS sample buffer supplemented with 10 mM DTT and boiled for 10 min to release bVAD-fmk-trapped proteins from the beads. The mixture was centrifuged for 1 min at 1000 *g*, and the soluble (streptavidin-bound) fraction was separated by gel electrophoresis alongside the unbound fraction.

### Enzyme-linked immunosorbent assay (ELISA)

TNF and IL-1β were measured using R&D ELISA kits following the manufacturers’ procedures. Total cytokine levels were calculated using GraphPad Prism by interpolating unknown sample values using a standard curve and multiplying by the sample dilution factor. Cytokine release was graphed using GraphPad Prism.

### Flow Cytometry

Flow cytometry was performed on total bone marrow, blood and spleen cells of 6-week-old male and female mice. Bones were flushed using a 27G needle attached to a syringe containing 5 mL PBS + 10% FBS and filtered through a 70 µM cell strainer. Spleens were mashed between two microscopy slides and filtered through a 70 µm cell strainer. Both total bone marrow cells and spleen cells were treated with 10 mL RBC lysis buffer, incubated for 10 min at room temperature and centrifuged for 5 min at 500 *g*. The pelleted bone marrow and spleen cells were resuspended and stained, and incubated on ice protected from light for up to 45 min. For the progenitor panel, bone marrow cells were stained for 45 minutes at 4°C in PBS containing the following fluorescently conjugated antibodies and dyes: FITC anti-mouse CD34, PE anti-mouse CD135, PE-Cy7 anti-mouse Sca-1, FVS700, BV420 anti-mouse CD16/32, APC-Cy7 anti-mouse CD117, BV510 anti-mouse CD45, APC Lineage for anti-mouse CD3a, anti-mouse CD11b, anti-mouse Gr-1 (Ly6G/Ly6C), anti-mouse B220 (CD45R) and anti-mouse Ter119. For the mature myeloid panel, bone marrow, blood and splenic cells were stained for 45 minutes at 4°C in PBS containing the following fluorescently conjugated antibodies and dyes: FITC anti-mouse C220 (CD45R), PE anti-mouse Ly6C, PE-Cy7 anti-mouse Ly6G, FVS700, APC anti-mouse CD115, APC-Cy7 anti-mouse CD11c and BV510 anti-mouse CD45.

Samples were acquired using a BD LSR Fortessa^TM^ X-20 Flow Cytometer, and unstained and single-stained samples were assayed in parallel to allow the appropriate voltages and compensation to be set. All data was analysed using FlowJo Software version 10.9.

### EdU pulse experiment

Click-iT Plus EdU Alexa Fluor 488 (ThermoFisher) was used and diluted to a stock concentration of 2 mg/mL EdU. 6-week-old WT and *Casp1.CDL* male mice were intraperitoneally injected with 100 µL of the 2 mg/mL EdU stock solution. 24 and 48 hours after EdU injection, blood from retro-orbital bleeds were collected in EDTA tubes, and femurs were harvested. Cell surface markers stained single cell solutions of bone marrow and total blood cells, incubated on ice protected from light for up to 45 min. Cells were washed with 1% BSA in PBS, spun down at 500 *g* for 5 min and stained for EdU using the manufacture’s Click-iT protocol. Briefly, cells were fixed using 100 µL Click-iT fixative (Component D), incubated for 15 min at room temperature, washed and permeabilised using 100µL 1x Click-iT saponin-based permeabilization buffer, incubated for 15 min at room temperature and then washed. Click-iT reaction buffer was added (PBS, CuSO4, Fluorescent dye AlexaF488, Reaction Buffer), incubated for 30 min at room temperature, and washed using 1x Click-iT saponin-based permeabilization buffer. The remaining staining of surface antigens with PE and PE-tandem antibodies were then performed, and incubated on ice protected from light for up to 45 min. The following fluorescently conjugated antibodies and dyes were used: AF488 EdU, BV421 anti-mouse CD11b, BV510 anti-mouse CD45, BV711 anti-mouse CD115, BV785 anti-mouse Ly6C, APC anti-mouse APC, APC-Cy7 anti-mouse CD117, live/dead marker FVS700, PE-Cy5 anti-mouse CD62L, PE-Cy7 anti-mouse Ly6G, BUV395 anti-mouse CD44. Samples were acquired using the BD LSR Fortessa^TM^ X-20 Flow Cytometer, and unstained and single-stained samples were assessed in parallel in order to set the appropriate voltages and compensation. All data was analysed using FlowJo Software version 10.9.

### Colony forming unit assay

Methylcellulose-based medium MethoCult (StemCell Technologies) was used to grow bone marrow cells for their differentiation into erythroid progenitor cells (MEP) and granulocyte-macrophage progenitor cells (GMP). Briefly, 3 x 10^5^ cells/mL were resuspended in IMDM media + 2% FBS + 1% Pen/Strep. 0.25 mL of cell suspension was mixed with 2.5 mL pre-warmed Methocult at 37°C. 1.1 mL of the Methocult-cell suspension mix was seeded per 35 mm^2^ round dish and incubated at 37°C + 5% CO_2_ for 10 days. At day 10, cells were lifted and the dish was twice washed with cold PBS to collect the remaining adherent cells. Cells were spun down at 500 *g* for 5 min, resuspended in PBS and stained, by incubating antibodies with cells on ice protected from light for up to 45 min using the following fluorescently conjugated antibodies: FITC anti-mouse CD34, PE anti-mouse CD135, PE-Cy7 anti-mouse Sca-1, FVS700, BV420 anti-mouse CD16/32, APC-Cy7 anti-mouse CD117, BV510 anti-mouse CD45, APC Lineage for anti-mouse CD3a, anti-mouse CD11b, anti-mouse Gr-1 (Ly6G/Ly6C), anti-mouse B220 (CD45R) and anti-mouse Terr119. Samples were acquired using a BD LSR Fortessa^TM^ X-20 Flow Cytometer, and unstained and single-stained samples were assayed in parallel in order to set the appropriate voltages and compensation. All data was analysed using FlowJo Software version 10.9.

### Data quantification and statistical analysis

All flow cytometry data were analysed using BD FlowJo X 10.07. Histology images were quantified using Fiji ImageJ. All statistical analyses were performed using GraphPad Prism 10. Data from pooled biological replicates were tested for normality using the Shapiro-Wilk test. Non-parametric data were further analysed by pairwise Mann-Whitney tests. Parametric datasets were analysed by one or two-way ANOVA with appropriate multiple testing correction (Tukey or Sidaks). Significance levels are indicated in each figure legend, with ns = p>0.05, * = p<0.05, ** = p<0.01, *** = p<0.001, **** = p<0.0001. Statistical analyses of each experiment are detailed in the corresponding figure legend.

