## Supplementary Figures for "Caspase-1 self-terminates protease activity to enforce homeostasis and prevent inflammasome-driven diseases"

### Supplementary Figures and Figure Legends

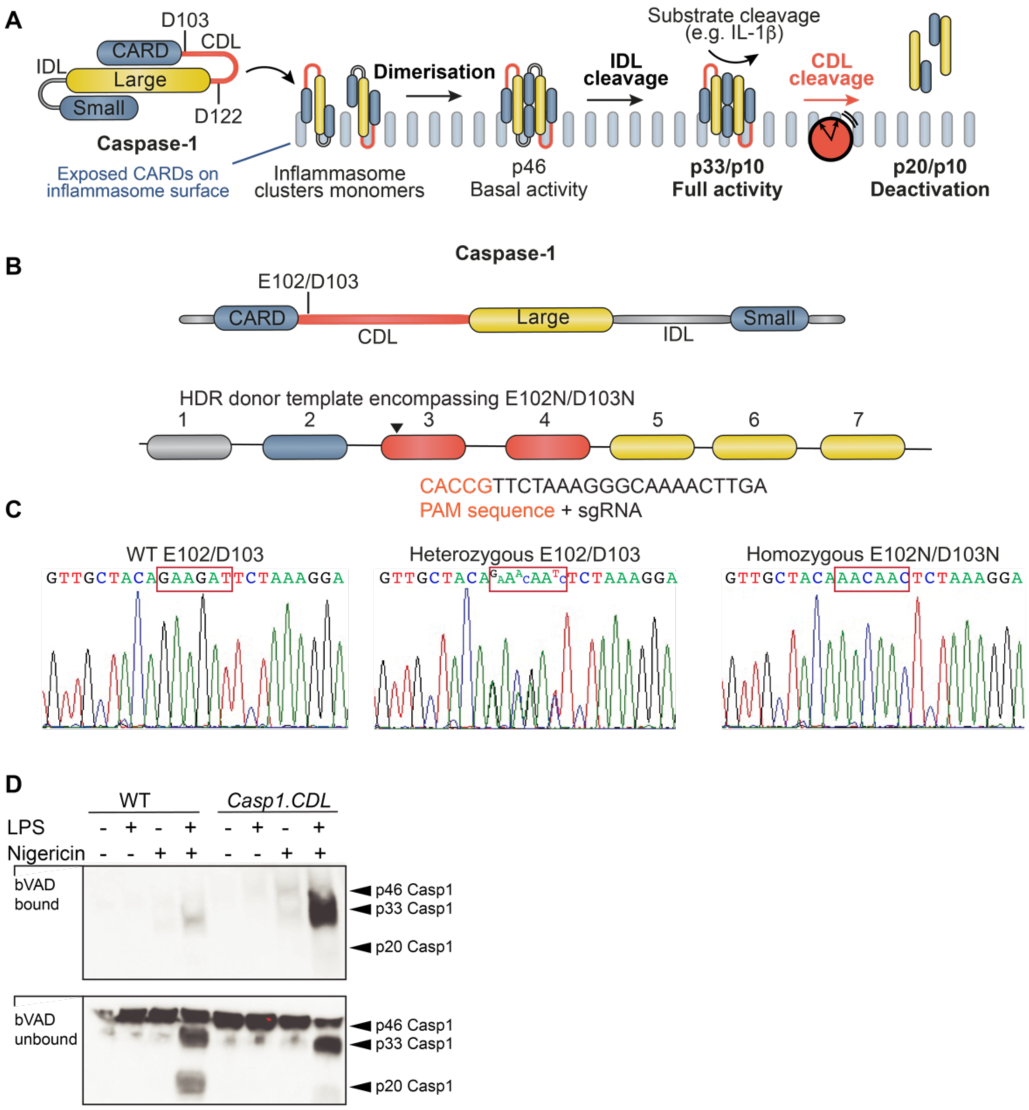


**Supplementary Figure 1. Generation of homozygous *Casp1.CDL* (E102N/D103N) mice.**
**a**, Schematic view of mechanisms of CASP1 activation and deactivation on the inflammasome, in which the CASP1 CARD domain linker (CDL) is labelled in red. **b-c**, Approach for gene knock-in by CRISPR/Cas9 gene editing. The HDR donor template encompassed the sequence ranging from exon 1 to exon 7 of the genomic sequence of CASP1 on Chr 9, and included a compound point mutation at E102N/D103N. The sgRNA sequence is shown. **c**, Sequencing by PCR screening of F1 mice. **d,** Pull-down of active CASP1 from WT versus *Casp1.CDL* BMDMs, using the biotin-VAD-fmk CASP1 activity probe (bVAD). BMDM were left untreated, or primed for 4 h with 100 ng/ml ultrapure *E. coli* LPS and then stimulated with nigericin for a further 1 h. The probe was applied to cells 0.5 h before nigericin. Streptavidin-coated beads pulled down active CASP1 bound to the biotin-labelled activity probe in mixed lysates/supernatants. Streptavidin-bound and unbound fractions were analysed by immunoblot using an antibody directed against the CASP1 large subunit. Representative blot of n=3 biological replicates. Related to Main Figure 1.

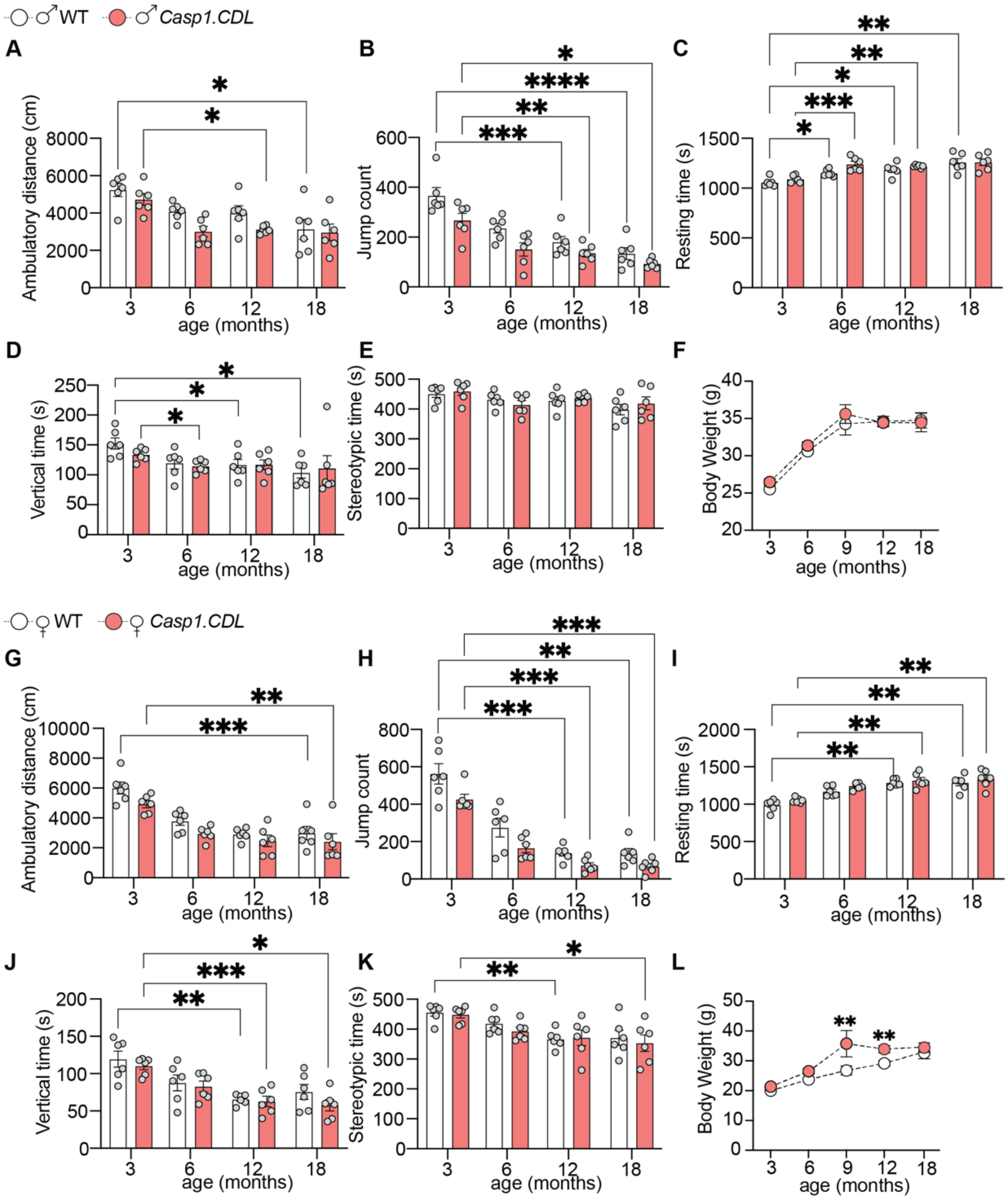


**Supplementary Figure 2. Age but not genotype affects the activity of WT and *Casp1.CDL* mice.**
**a-l**, WT and *Casp1.CDL* male (a-f) and female (g-l) mice were aged, and assessed for activity and body weight at 3-, 6-, 12- and 18 months of age. **a-e+g-k**, open field test with quantification of ambulatory distance (a+g), Jump count (b+h), resting time (c+i), vertical time (d+j) and stereotypic time (e+k). **f+l**, body weight. All behaviour data is represented as mean ± SEM with dots representing individual mice (n=6 mice for each sex and age group, quantified from 2-3 independent biological experiments). Data were verified for normality using a Shapiro-Wilk test, and analysed by two-way ANOVA with Tukey's multiple testing correction. Statistical significance: * $p$≤0.05; ** $p$≤0.01, *** $p$≤0.001, **** $p$≤0.0001. Related to Main Figure 2.

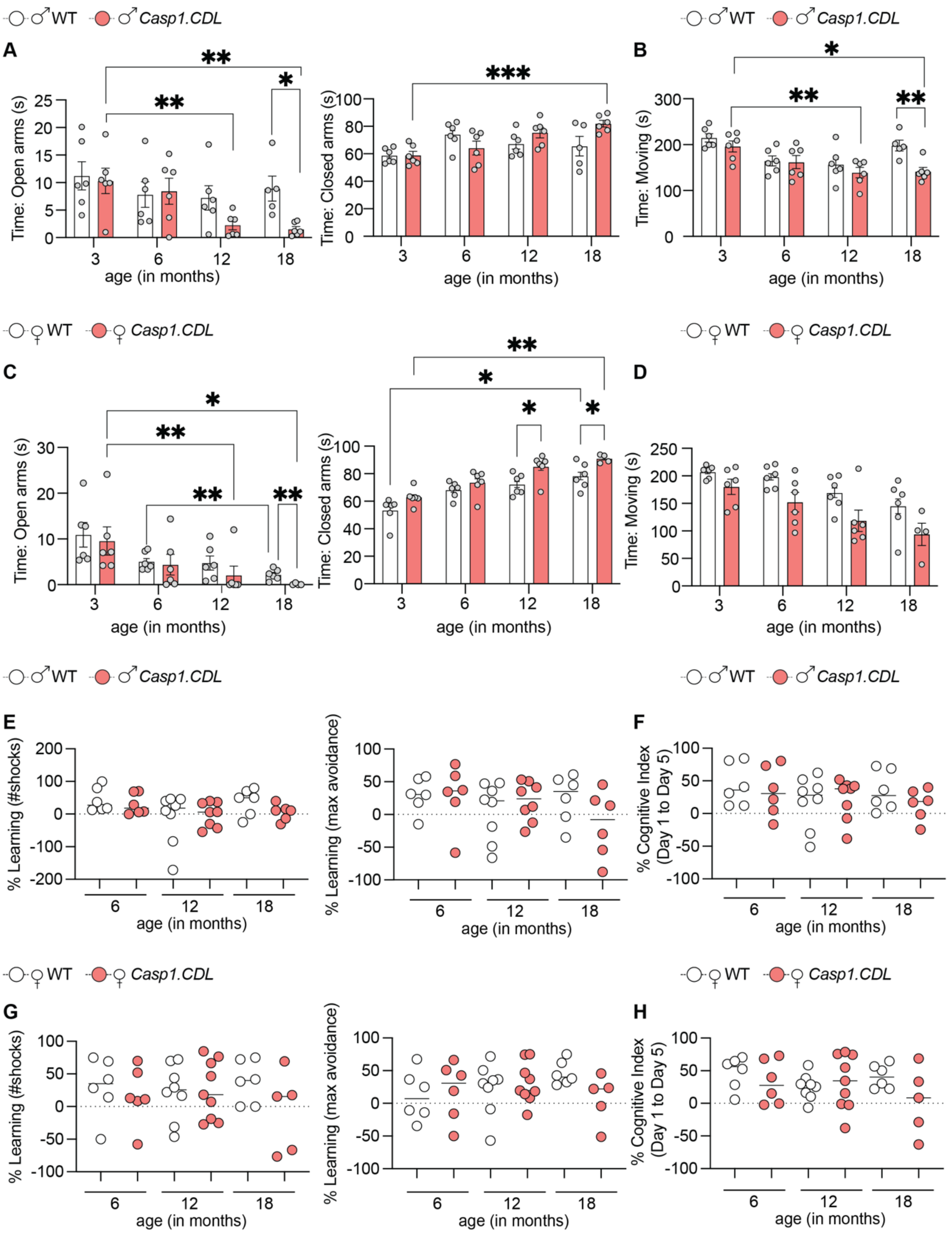


**Supplementary Figure 3. CASP1 CDL mutation drives steady-state anxiety-like behaviour in male and female mice.**

WT and *Casp1.CDL* male (a-b, e-f) and female (c-d, g-h) mice were aged to 18 months and assessed for behaviour. **a-d**, Mice of 6-, 12- and 18 months of both sexes were tested in the elevated plus maze (EPM), and assessed for time spent in open and closed arms (a, c) and movement over the duration of the test (b, d). **e-h**, Active place avoidance (APA) test was performed for 5 constitutive days with 6-, 12- and 18-month-old mice of both sexes. % Learning between Day 1 and Day 5 was quantified, and shown as % learning based on shock number, % learning based on maximal time to avoid a shock and a multi-parameter indicator of cognitive function (% cognitive index). Each dot represents individual mice (n=6-8 male and n=6-8 female mice tested in 2-3 independent cohorts). Data were verified for normality using the Shapiro-Wilk test, and analysed by two-way ANOVA with Tukey's multiple testing correction. Statistical significance: * $p \leq 0.05$; ** $p \leq 0.01$, *** $p \leq 0.001$, **** $p \leq 0.0001$. Related to Main Figure 2.

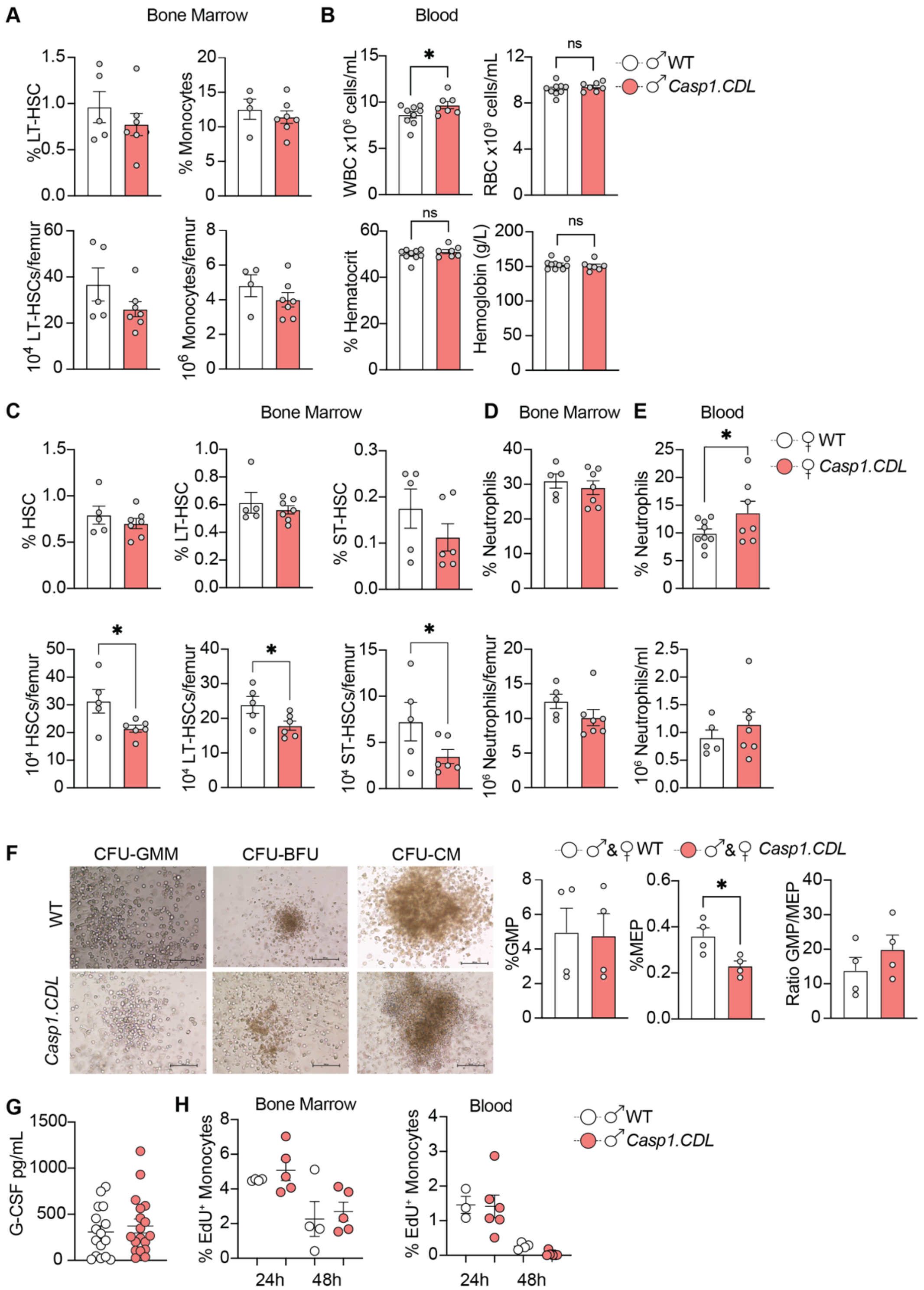


**Supplementary Figure 4. *Casp1.CDL* mice exhibit enhanced steady-state granulopoiesis.**
**a-e,** 6-week-old male (a-b) and female (c-e) mice were assessed for: percentage and total counts of bone marrow hematopoietic stem cells (a, c); mature neutrophils and monocytes in bone marrow (a, d) and blood (e); and total circulating white blood cells (WBC), red blood cells (RBC), haematocrit and haemoglobin. **f,** Bone marrow cells were cultured in methylcellulose-based medium for their differentiation into erythroid progenitor cells (MEP) and granulocyte-macrophage progenitor cells (GMP). Images are representative of cells at day 10 in culture (scale bar 100 µm). Cell differentiation into MEPs and GMPs was assessed by flow cytometry. **g**, Quantification of serum G-CSF in 6-week-old male and female mice. **h,** 6-week-old male mice were intraperitoneally injected with 2 mg/mL EdU. The percentage of $EdU^+$ monocytes in the bone marrow and the blood was analysed 24 h and 48 h post-injection. **a-h**, Scatterplot dots represent data for individual mice. Data are n=5-6 mice/genotype from 3-4 independent biological experiments (a-e), n=4 mice/genotype from 4 independent biological experiments (f), serum from n=16-18 male and female mice/genotype from 4-5 independent experiments (g), and n=3-5 mice/genotype from 2 independent biological experiments (h). Data were verified as non-parametric using the Shapiro-Wilk test, and analysed by the Mann-Whitney U-test. Statistical significance: * $p \le 0.05$; ** $p \le 0.01$, *** $p \le 0.01$. Related to Main Figure 3.

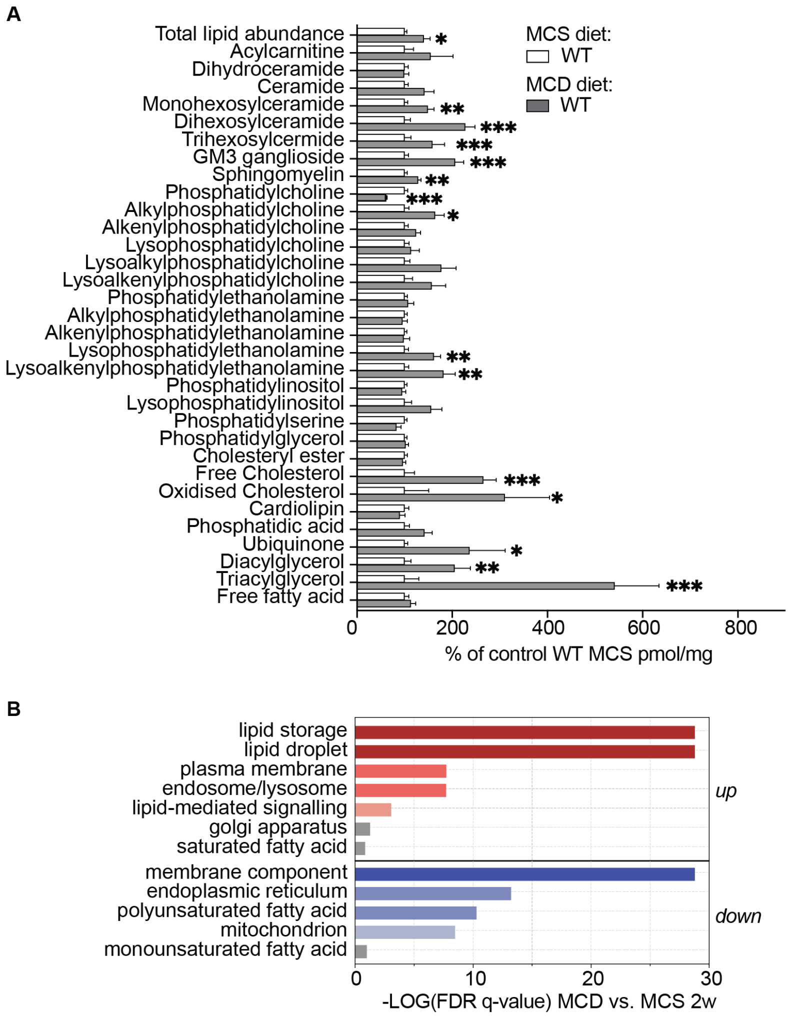


**Supplementary Figure 5. The MCD diet alters hepatic lipid abundance.**

**a** Hepatic lipid abundance of the measured lipid classes and subclasses of MCD-fed WT mice (grey bars) were compared with the MCS-fed WT mice (white bars). Lipid abundance in MCD-fed mice is presented as a percentage relative to the MCS control. Data are mean ± SEM from n=8 mice per group. Data were verified as non-parametric using the Shapiro-Wilk test, and analysed by Multiple Mann-Whitney U-test. Statistical significance: *p < 0.05, **p < 0.01, ***p < 0.001. **b,** Lipids significantly up- or down-regulated in MCD-fed WT diet mice were analysed using lipid ontology enrichment for function, cellular component and chemical and physical properties. Related to Main Figure 4.

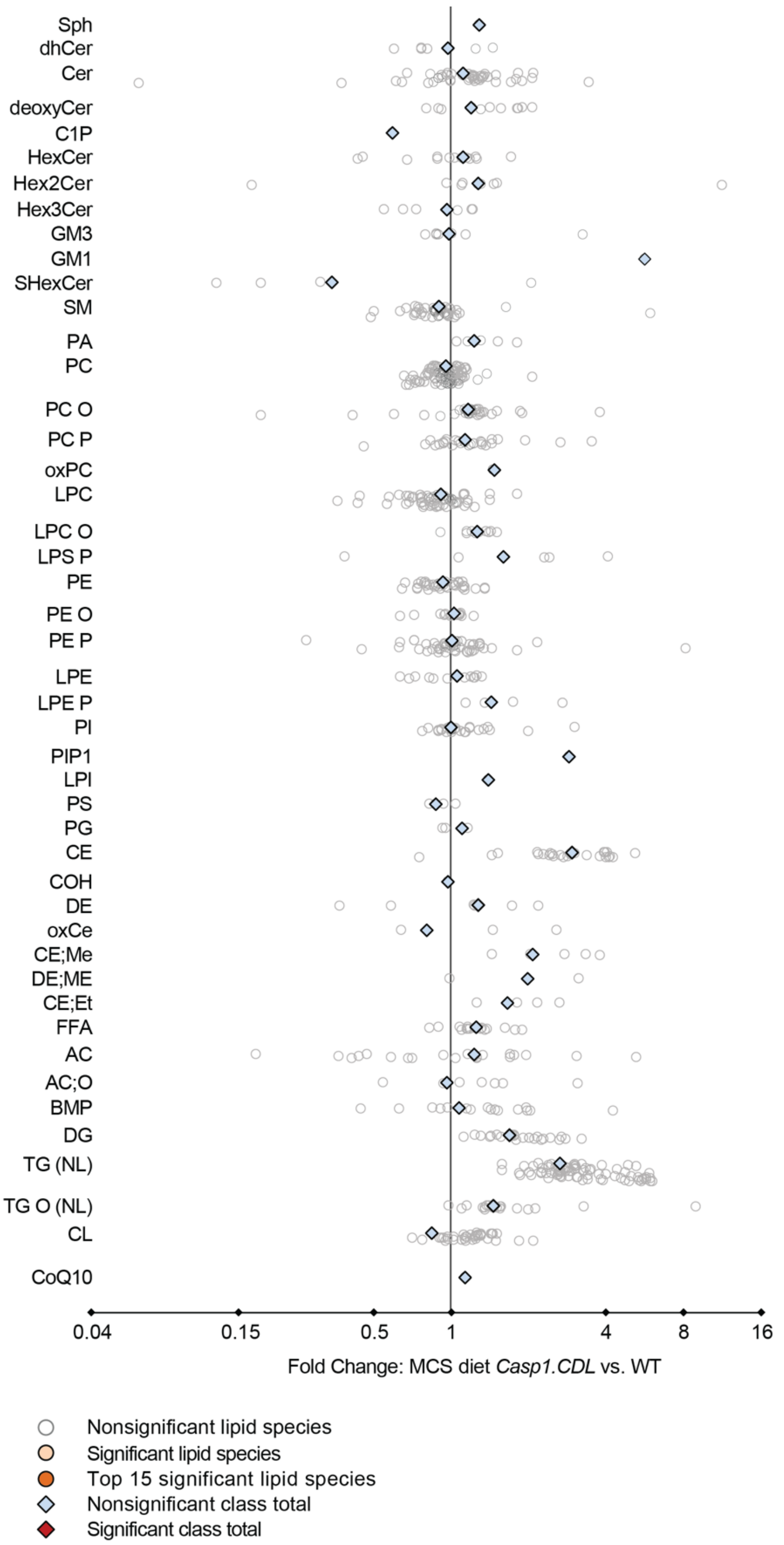


**Supplementary Figure 6. CASP1 CDL mutation did not affect hepatic lipid abundance in MCS-fed mice.** Forest plot showing the fold changes in hepatic lipid species and classes between MCS-fed *Casp1.CDL* mice compared to MCS-fed WT mice. Data are fold change with upper and lower confidence intervals (n=8 WT, n=9 *Casp1.CDL* mice). A paired t-test with false discovery rate testing correction identified no statistically significant differences. Related to Main Figure 4.

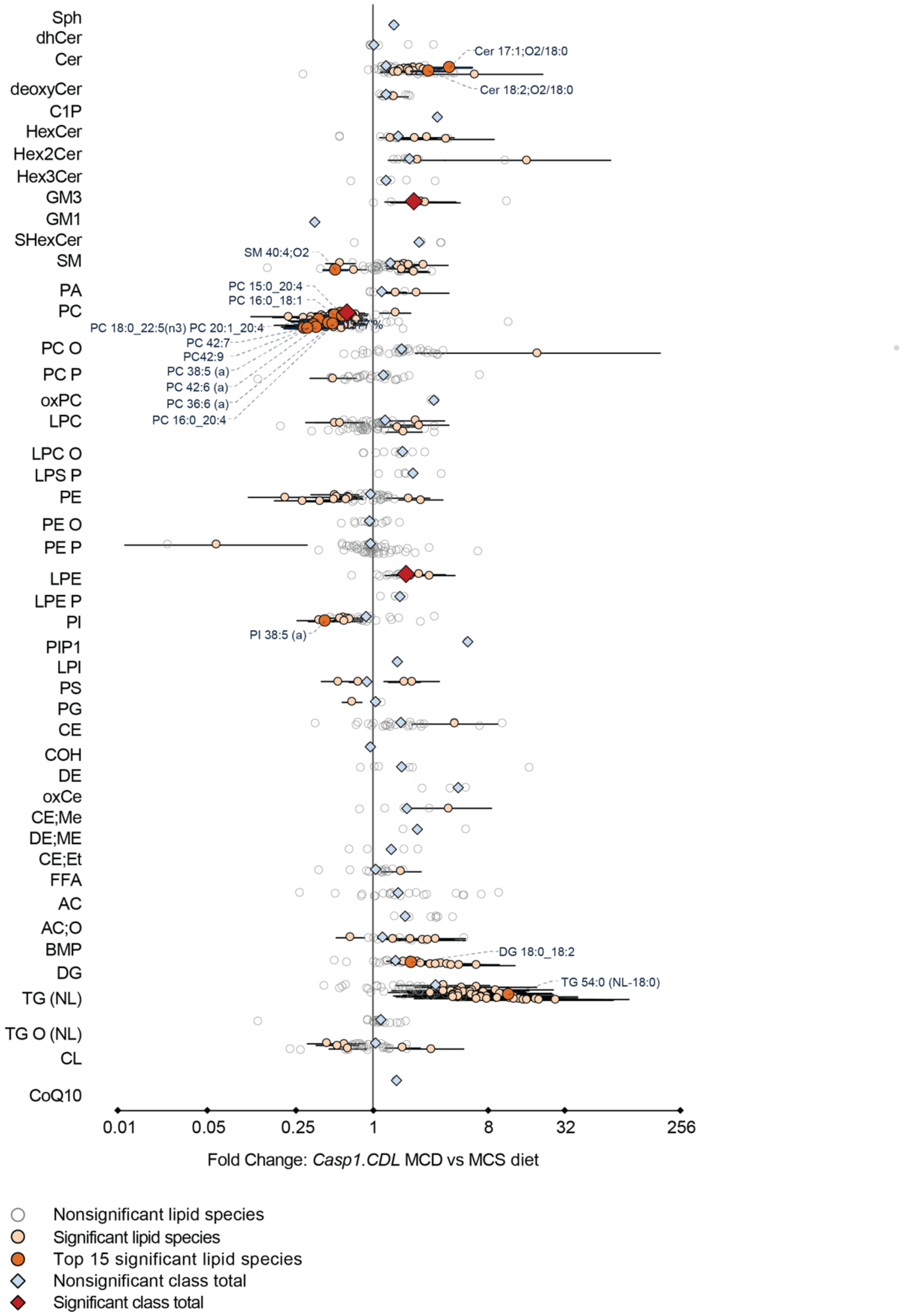


**Supplementary Figure 7. The MCD diet altered hepatic lipid abundance in *Casp1.CDL* mice.**
Forest plot showing the fold changes in hepatic lipid species and classes between MCD-fed *Casp1.CDL* mice compared to MCS-fed *Casp1.CDL* mice. Data are fold change with upper and lower confidence intervals (n=9 MCS-fed, n=10 MCD-fed mice). A paired t-test with false discovery rate testing correction identified statistically significant differences. Related to Main Figure 4.

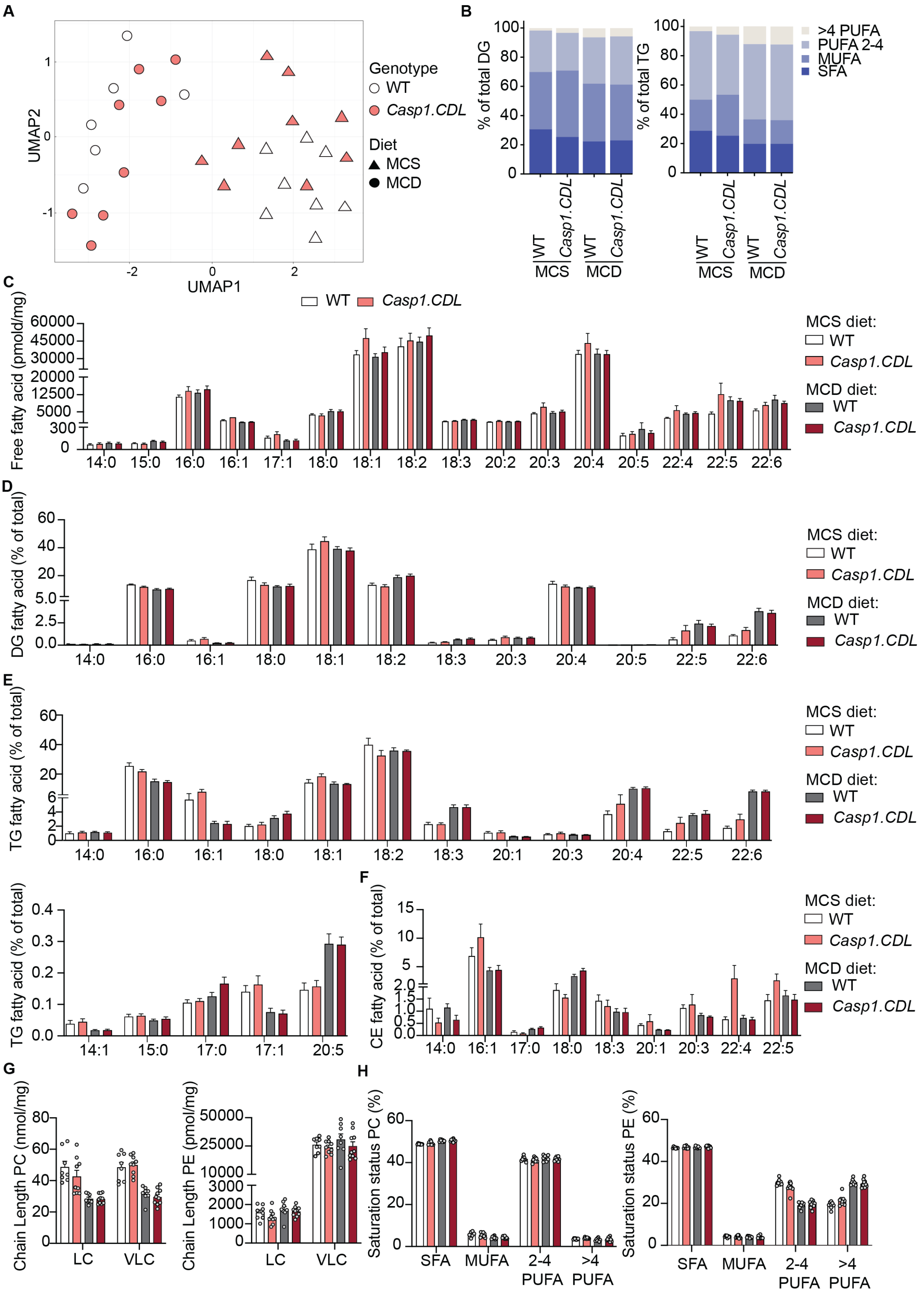


**Supplementary Figure 8. MCD diet, but not CASP1 CDL mutation, markedly affected hepatic lipid abundance.** **a,** UMAP plot shows the distribution of the hepatic lipidome of MCD-fed WT (white circles), MCD-fed *Casp1.CDL* (pink circles), MCS-fed WT (white triangles) and MCS-fed *Casp1.CDL* (pink triangles) mice. **b,** Fatty acid (FA) composition of hepatic diglycerides (DG) and triglycerides (TG). **c**, Hepatic free fatty acid (FFA) levels. **d-f**, FAs comprising hepatic DG, TG and cholesterol esters (CE), shown as the percentage of each lipid class. Histograms are mean ± SEM n=8-10 mice/group. Data were verified as non-parametric using the Shapiro-Wilk test, and Mann-Whitney U-test with Holm-Šídák correction identified no significant differences between MCD-fed genotypes. Related to Main Figure 4.

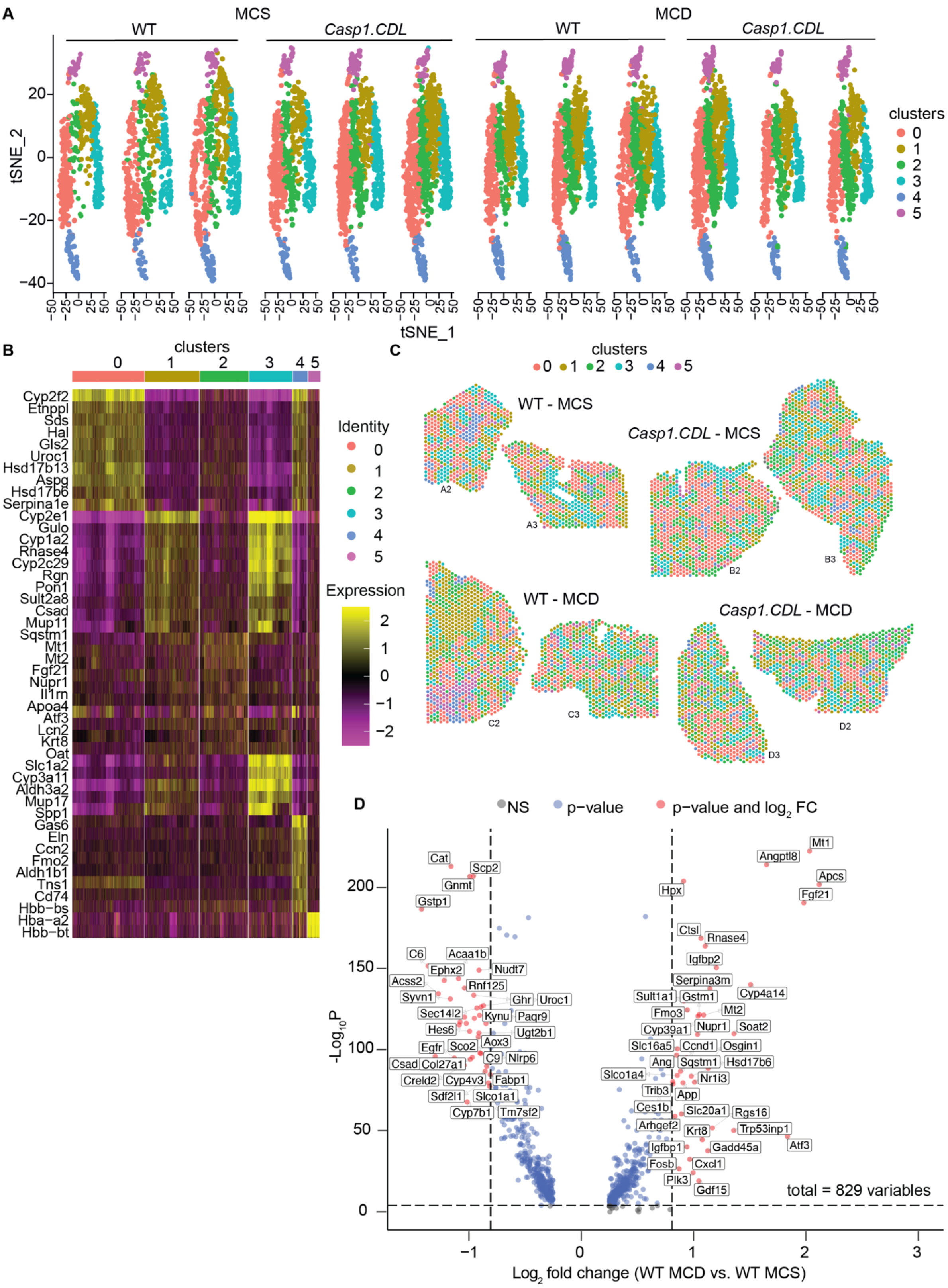


**Supplementary Figure 9. 10xGenomics spatial transcriptomics identifies six conserved spot clusters present in all liver samples, regardless of genotype or diet.**
Spatial mapping of transcriptomic clusters (visium spots) in mice fed a methionine-choline supplemented or deficient (MCS, MCD) diet for 2 weeks. 55µm diameter spots were clustered using principal components derived from their transcriptomic signatures (labelled as cluster 0-5), and visualised in a 2-dimensional t-SNE scatter plot across all liver samples (**a**). **b,** Heatmap showing the top 10 differentially expressed genes per cluster. **c,** spatial x-y coordinates of spots belonging to each cluster on the remaining liver sections that were not shown in Figure 4b. **d,** Volcano plot shows genes of spot cluster 0 that were differentially expressed in MCD-fed versus MCS-fed WT mice ($log_2FC > 0.75$ and FDR = 0.05). Statistical analysis was performed using a two-sided Wilcoxon rank-sum test with Bonferroni multiple testing correction. **a-d,** Data are from n=3 mice per experimental group. Related to Main Figure 5.

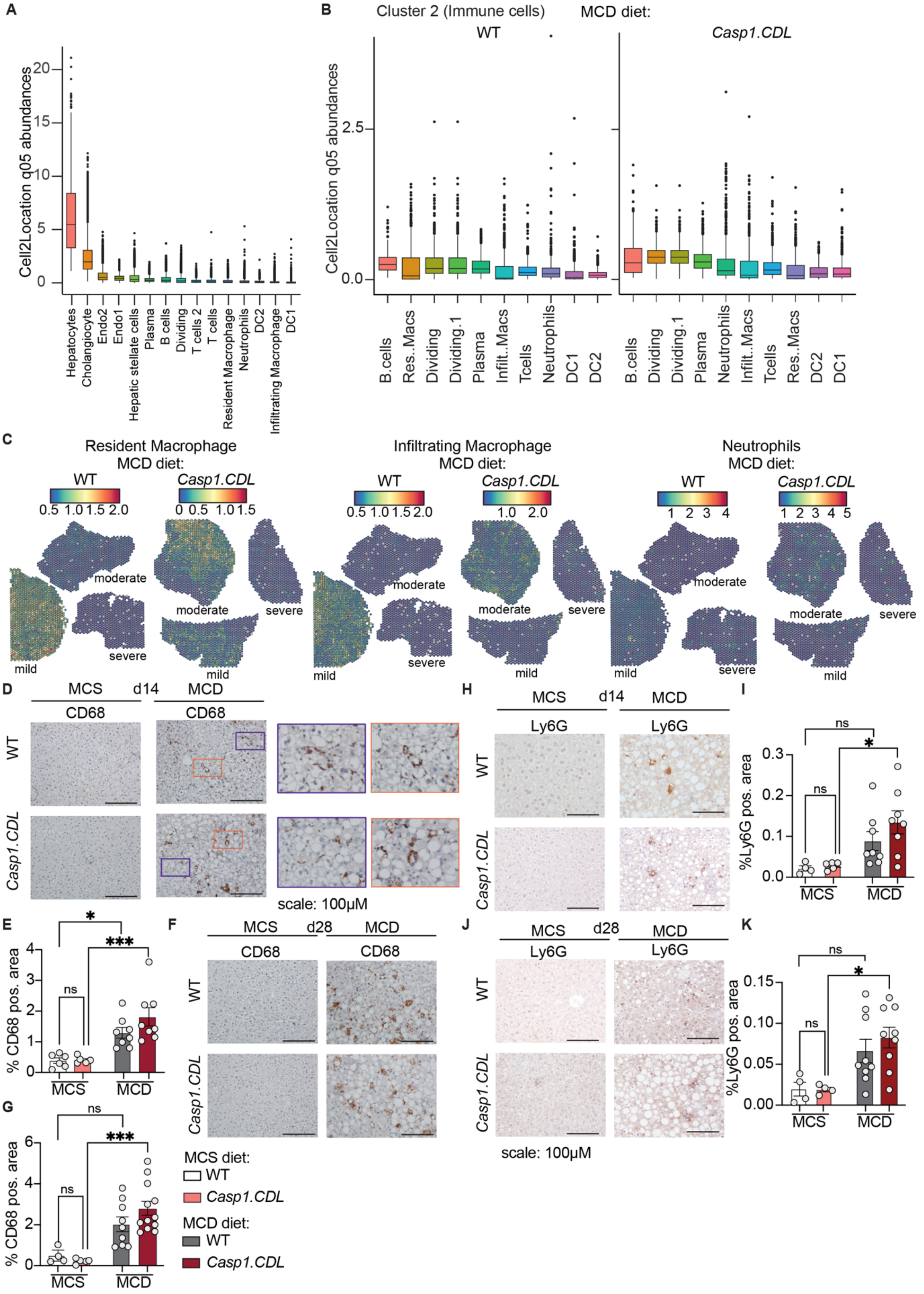


**Supplementary Figure 10. The MCD diet induces hepatic immune responses, including macrophage and neutrophil infiltration in *Casp1.CDL* mice.**

**a-b,** Box and whisker plot of predicted abundances of all cell types (as re-analysed from GSE129516) captured in cluster 2 spots. Box bounds and whiskers show interquartile range (IQR) and extreme values (within 1.5x the IQR), while outliers are shown as data points outside the whiskers. **c**, Cell2location predicted the abundance of resident and infiltrating macrophages and neutrophils as re-analysed from GSE129516 in visium spots, and plotted as spatial features on liver sections. For each condition, liver sections with mild (less than 20% steatosis), moderate (20-50%

steatosis) and severe (above 50% steatosis) pathology were analysed. **d-k**, DAB immunohistochemical staining of livers for mice fed an MCD or control MCS diet for 2 weeks (d, e, h, i) or 4 weeks (f, g, j, k). Livers were stained for macrophages (Kupffer cells, infiltrating macrophages) using anti-CD68 and neutrophils using anti-Ly6G. CD68+ cells exhibit stark differences in morphology, with these cells either elongated (highlighted in the enlarged purple boxes, d) or round (highlighted in the enlarged orange boxes, d). CD68 and Ly6G positive area was quantified using Fiji (ImageJ) analysis of the average signal of three liver pieces per mouse. Scale bar 100 µm. **e, g, i, k,** Scatterplots show mean ± SEM, including data for each mouse (dots) from n=4-9 mice/condition from 4-5 independent experiments. Data were verified for normality using the Shapiro-Wilk test, and analysed by one-way ANOVA. Statistical significance: * *p*≤0.05; ** *p*≤0.01, *** *p*≤0.001, **** *p*≤0.0001. Related to Main Figure 6.

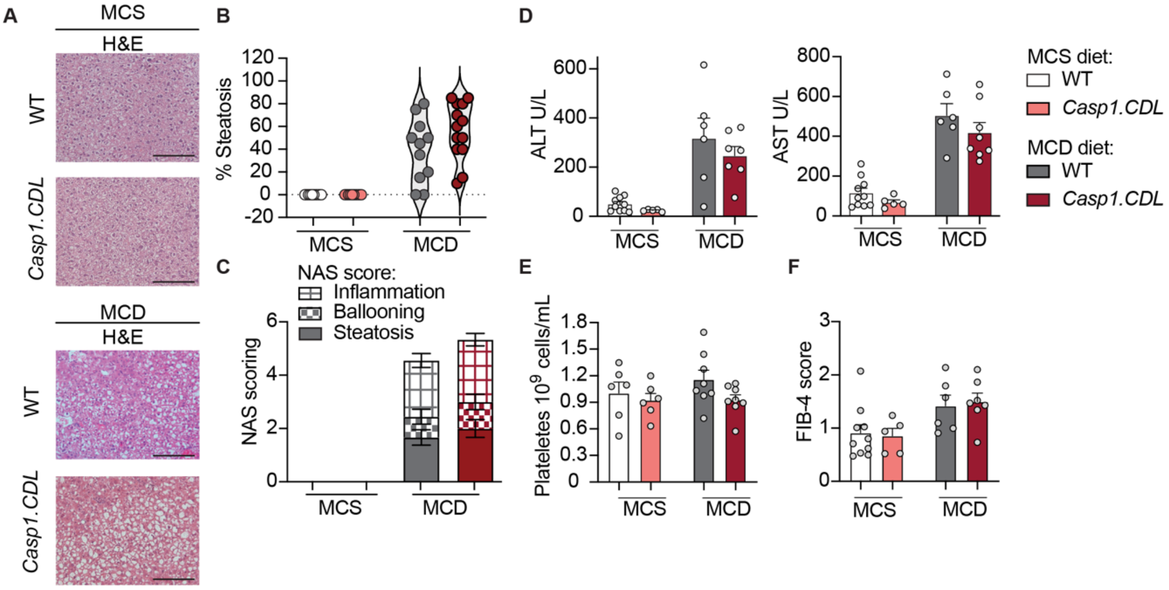


**Supplementary Figure 11. WT and *Casp1*.CDL show similar liver pathology and serum markers of liver damage after 4 weeks of MCD feeding to model advanced MASLD.**

12-week-old male mice were fed a methionine-choline supplemented diet (MCS) or a methionine-choline deficient diet (MCD) for 4 weeks and assessed for indicators of liver pathology. **a,** representative images for liver H&E staining (scale bar 100 µM). **b-c**, H&E-stained sections were blindly assessed for liver pathology according to percentage of steatosis (b), and scored using the NAS score (an additive score of steatosis, hepatocyte ballooning and inflammation, c). **d,** Serum levels of liver enzymes, alanine transaminase (ALT) and aspartate aminotransferase (AST), were measured as indicators of liver damage. **e,** The abundance of platelets in whole blood. **f,** FIB-4 score was calculated from ALT, AST and platelet counts. **c-f**, Dots in violin plots and scatterplots are data from individual mice (n=5-10 mice/group from 3-4 independent experiments), except serum analyses, where two mice/group were pooled for analyte detection (d). Data are mean ± SEM, were verified for normality using the Shapiro-Wilk test, and analysed by one-way ANOVA, which identified no significant differences between MCD-fed genotypes. Related to Main Figure 7.

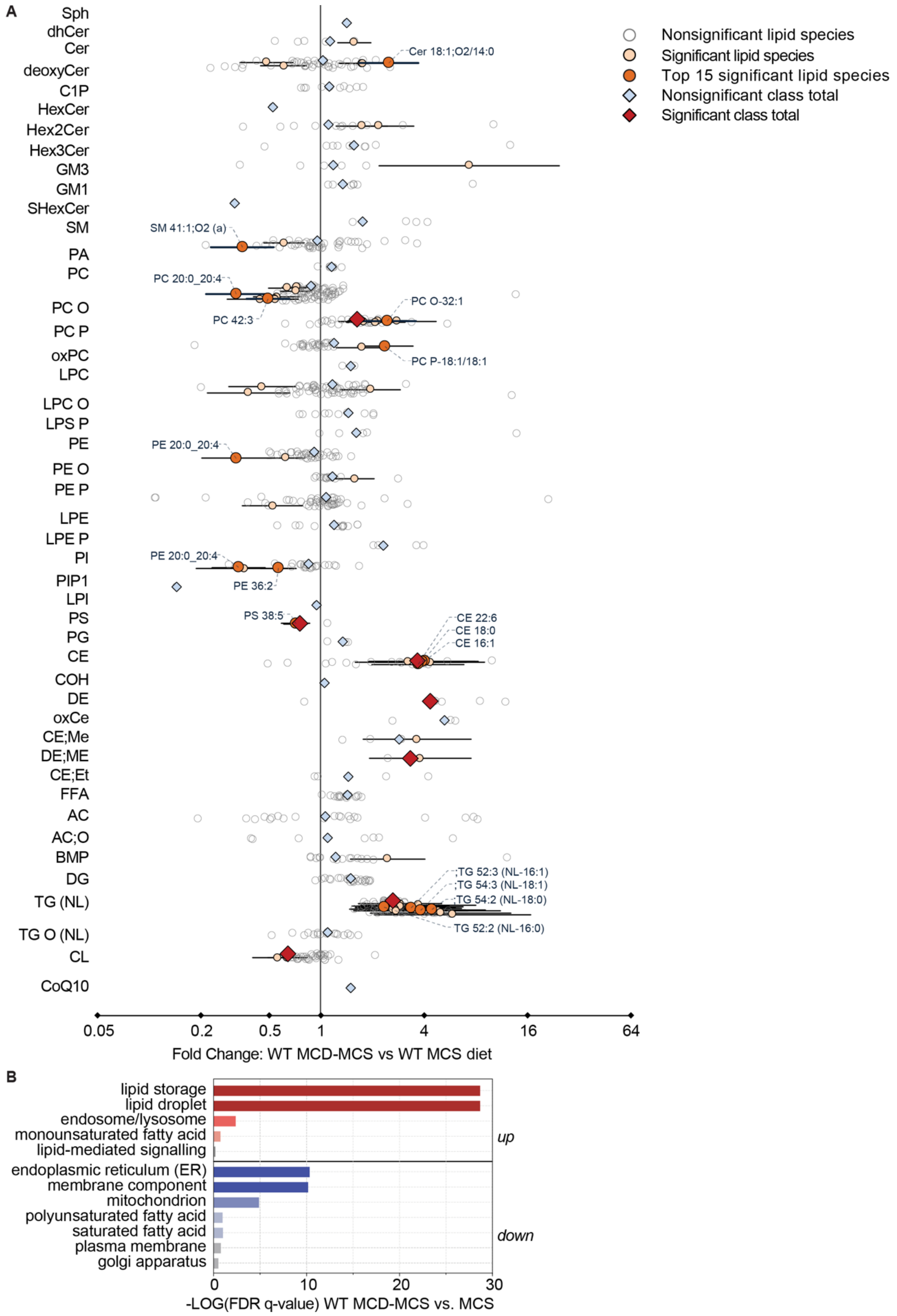


**Supplementary Figure 12. MCD-induced hepatic lipid accumulation is ameliorated after MCS feeding in WT mice.**

WT mice were fed an MCD diet for 4 weeks to induce liver damage, and then switched to the control MCS diet for 7 days to allow liver healing (MCD-MCS), or fed the MCS diet for 5 weeks (MCS)**. a**, Forest plot showing the fold change difference in hepatic lipid species and classes. **b**, Lipid ontology enrichment for function, cellular component and chemical and physical properties that were significantly up or down-regulated in MCD-MCS versus MCS-fed mice. Data are fold change with upper and lower confidence intervals (n= 7 WT MCS and n=9 WT MCD-MCS-fed mice). A paired t-test with false discovery rate testing correction identified statistically significant differences. Statistical significance: * *p*≤0.05; ** *p*≤0.01, *** *p*≤0.01. Related to Main Figure 7.

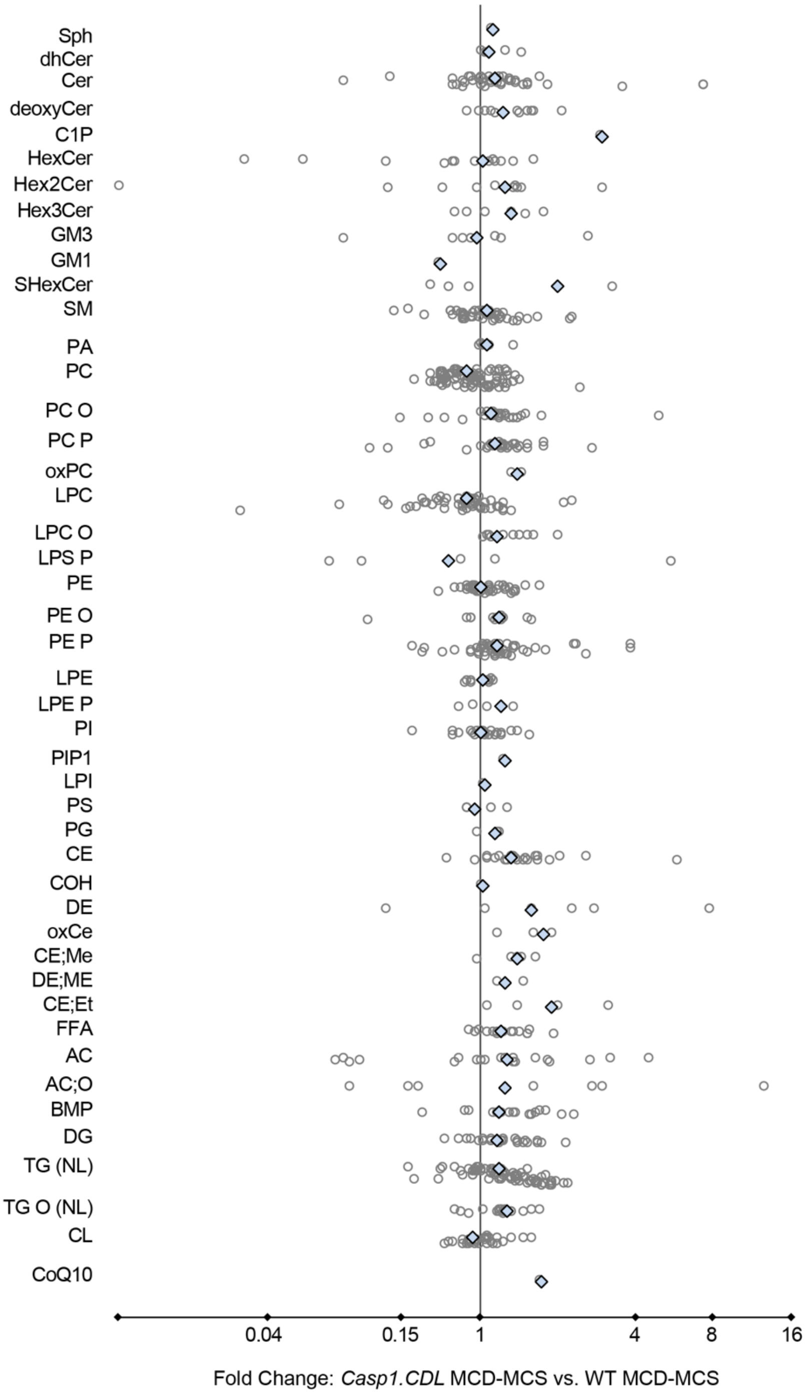


**Supplementary Figure 13. CASP1 CDL mutation did not affect the return to homeostatic levels of hepatic lipids in mice healing from MCD-induced liver damage.**

Mice were fed an MCD diet for 4 weeks to induce liver damage, and then switched to the control MCS diet for 7 days to allow liver healing (MCD-MCS)**.** Forest plot showing the fold change difference in hepatic lipid species and classes in *Casp1.CDL* versus WT MCD-MCS-fed mice. Data are fold change with upper and lower confidence intervals (n=9 WT, n=11 *Casp1.CDL* MCD-MCS-fed mice). A paired t-test with false discovery rate testing correction identified no statistically significant differences between the genotypes. Related to Main Figure 7.

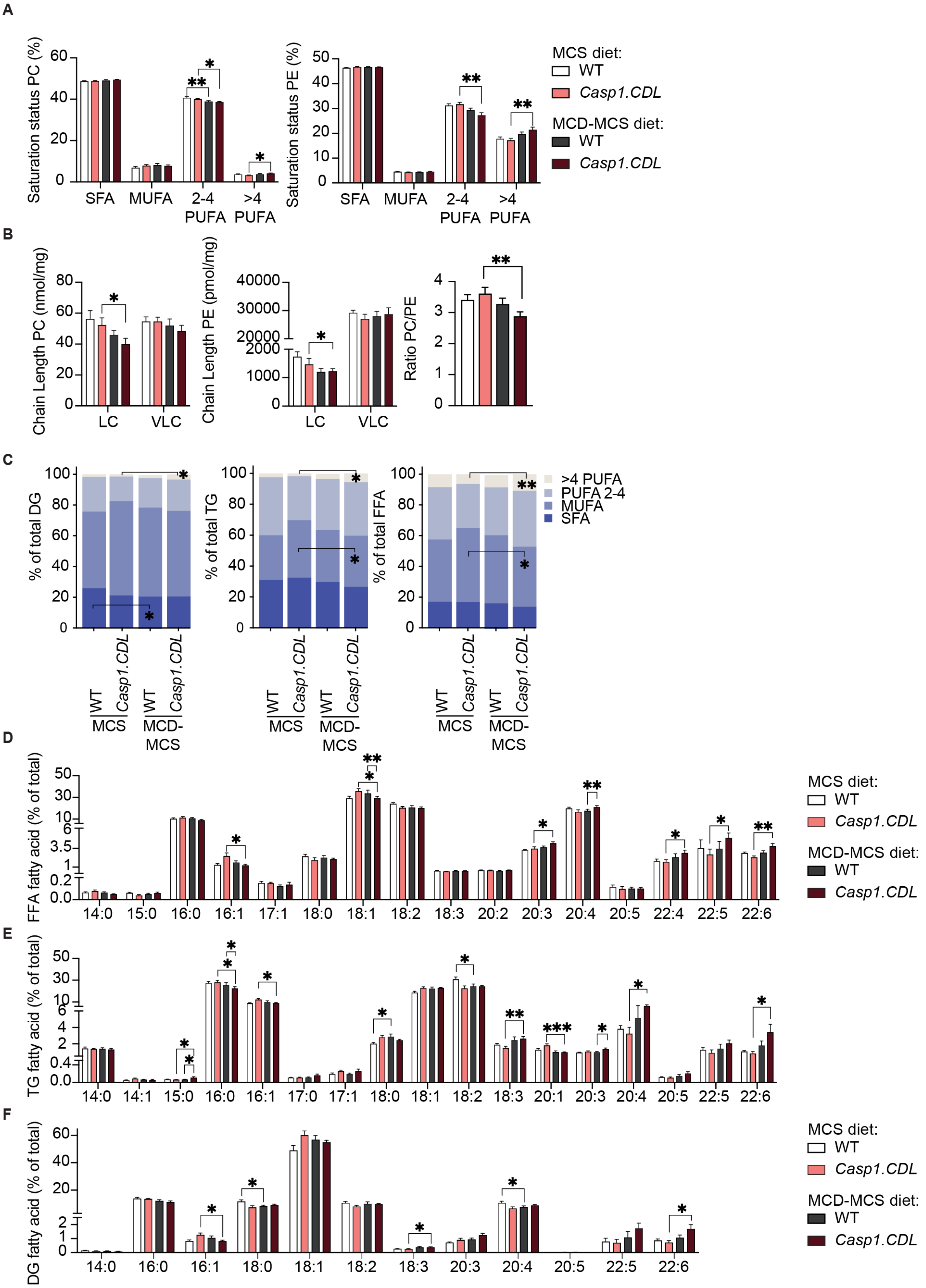


**Supplementary Figure 14. CASP1 CDL mutation suppressed the return to homeostatic proportions of individual lipid classes in mice healing from MCD-induced liver damage.**

*Casp1.CDL* versus WT mice were fed an MCD diet for 4 weeks to induce liver damage, and then switched to the control MCS diet for 7 days to allow liver healing (MCD-MCS), and compared to MCS-fed mice. **a-b**, Chain length and saturation status of fatty acid composition of hepatic phospholipids, phosphatidylcholine (PC) and phosphatidylethanolamine (PE). **c,** Fatty acid (FA) composition of hepatic diglycerides (DG), triglycerides (TG) and total fatty acids. **d**, Hepatic free fatty acid (FFA) levels. **e-f**, FAs comprising hepatic triglycerides (TG) and diglycerides

(DG), shown as the percentage of each lipid class. Histograms are mean ± SEM (n=7 MCS-fed WT; n=9 for MCD-MCS-fed WT; n=8 for MCS-fed *Casp1.CDL*; n=11 for MCD-MCS-fed *Casp1.CDL* mice). Data were assessed for normality using the Shapiro-Wilk test. Non-parametric data was assessed by Mann Whitney U-test with Holm-Šídák correction (a-c) and parametric data was assessed by two-way ANOVA with Tukey's multiple testing correction (d-f). Statistical significance: * $p \leq 0.05$; ** $p \leq 0.01$. Related to Main Figure 7.
